# Multi-omics Analysis Identifies IgG2b Class-Switching with ALCAM-CD6 Co-Stimulation in Lymph Nodes During Advanced Inflammatory-Erosive Arthritis

**DOI:** 10.1101/2022.10.27.514103

**Authors:** H. Mark Kenney, Javier Rangel-Moreno, Yue Peng, Kiana L. Chen, Jennifer Bruno, Abdul Embong, Elizabeth Pritchett, Jeffrey I. Fox, Sally Quataert, Gowrishankar Muthukrishnan, Ronald W. Wood, Benjamin D. Korman, Jennifer H. Anolik, Lianping Xing, Christopher T. Ritchlin, Edward M. Schwarz, Chia-Lung Wu

## Abstract

Defective lymphatic drainage and B-cell translocation into joint-draining lymph node sinuses are pathogenic phenomena in patients with severe rheumatoid arthritis (RA). However, the molecular mechanisms underlying this lymphatic dysfunction remain poorly understood. Here, by utilizing spatial and single-cell transcriptomics in tumor necrosis factor transgenic (TNF-Tg) mice, we characterized functional genomic changes in popliteal lymph nodes (PLNs) of “Early” and “Advanced” RA to determine the mechanisms orchestrating B-cell differentiation. We first show that *Ighg2b* expression localized to Marco^+^ sinuses and negatively correlated with bone volumes in ipsilateral joints. We further reveal that Advanced PLNs exhibited a concomitant accumulation of iron-laden macrophages and T-cells. Mechanistically, crosstalk between ALCAM^+^ macrophages and CD6^+^ T-cells was identified as a co-stimulatory pathway promoting IgG2b class-switching. These findings were validated by immunohistochemistry, flow cytometry, ELISPOT, and clinical correlates. Collectively, we propose that ALCAM-CD6 co-stimulation activates T cells, initiating IgG2b class-switching and plasma cell differentiation in RA flare.

## Introduction

Rheumatoid arthritis (RA) is a complex disease driven by both chronic innate immune pathology (e.g. overproduction of tumor necrosis factor; TNF) that most prominently contributes to joint inflammation, pain and connective tissue destruction, and autoimmune features that include diagnostic autoantibodies (e.g. anti-citrullinated protein antibodies; ACPAs) (Firestein, 2014) and B/T-cell dysregulation (Meednu et al., 2022; Scheel et al., 2011). Among the enigmas of RA pathogenesis, how chronic “sterile” inflammation in affected joints contributes to the development of autoantibodies remains an important open question. It is also known how RA progression involves lymphatic dysfunction and alterations in joint draining lymph nodes (Bouta et al., 2018). Specifically, RA patients exhibit reduced lymphatic drainage from inflamed joints (Bell et al., 2020; Bouta et al., 2017), and power doppler ultrasound studies (PDUS) also demonstrate dynamic intra-parenchymal changes and vascular flow modulation of draining lymph nodes linked to the underlying pathologic process (Manzo et al., 2016; Manzo et al., 2011). Interestingly, RA joint-draining lymph nodes contain large numbers of quiescent, polyclonal CD23^+^/CD21^hi^ B-cells in inflamed nodes (Bin cells), whose pathogenic role remains unknown (Keffer et al., 1991; Kuzin et al., 2016).

To elucidate the aforementioned RA pathology and developing effective interventions, we characterized the natural history of inflammatory-erosive arthritis in TNF-transgenic (Tg) mice, which is a well-established autoantibody independent model of RA (Keffer et al., 1991). Interestingly, the 3647 TNF-Tg mouse line, which contains one copy of the transgene, recapitulates all the draining lymph node features and lymphatic dysfunction observed in RA patients (Bouta et al., 2018). During “Early” arthritis, the ankle draining popliteal lymph nodes (PLNs) demonstrate marked volume expansion and increased blood flow (Bouta et al., 2018). PLN expansion persists to ∼8-months of age in male TNF-Tg mice, followed by stochastic and asymmetric collapse of PLNs associated with reduced volume and blood flow, and rapid onset of “Advanced” arthritis with severe synovitis and bone loss (Bouta et al., 2018). Of note, female TNF-Tg mice die from cardiopulmonary disease by ∼5 months of age (Bell et al., 2019), which limits the potential investigation of the chronic lymphatic phenotypes and mechanisms.

Histologic assessments of PLNs from TNF-Tg mice with Early and Advanced arthritis provide insight into the cellular mechanisms mediating these dynamic lymph node changes (Bouta et al., 2018; Li et al., 2010; Li et al., 2011). In Early arthritis, the CD23^+^/CD21^hi^ Bin cells accumulate in PLN follicles while afferent CXCL13-expressing macrophages egress through the expanded sinuses. Collapsed PLNs in Advanced arthritis are characterized by translocated Bin cells into the sinuses (Bouta et al., 2018; Li et al., 2010; Li et al., 2011), presumably via a CXCL13 gradient produced by stagnant macrophages that cannot egress due to the lack of lymphatic vessel contractions and passive flow (Bouta et al., 2018). This Bin cell “clogging” of efferent lymphatics is hypothesized to exacerbate inflammatory-erosive arthritis, as their removal with anti-CD20 B-cell depletion therapy in TNF-Tg mice restores lymphatic flow and ameliorates Advanced arthritis (Bouta et al., 2018; Li et al., 2013; Li et al., 2010).

Recent attempts to elucidate the mechanisms responsible for lymphatic dysfunction and Bin cell accumulation and translocation in PLN sinuses during Early vs Advanced arthritis in TNF-Tg mice utilized various bulk-tissue approaches (i.e., bulk genomics, flow cytometry and *ex vivo* cultures), which failed to identify molecular changes related to arthritic severity (Bouta et al., 2018). Unfortunately, these experiments were limited by contamination of follicular B-cells with the inability to differentially assess B-cell phenotypes based on spatial location, such as those within the PLN sinuses. However, spatial and single-cell transcriptomic technologies have overcome these limitations and transformed several fields, including RA etiology and pathogenesis (Bhamidipat and Wei, 2022; Wei et al., 2020) Thus, we aimed to utilize these approaches to 1) examine if arthritic severity of afferent ankle joints correlates with unique phenotypes of B-cells within PLN sinuses of TNF-Tg mice and 2) identify potential co-stimulatory pathways between hematopoietic cells that could contribute to the initiation of PLN immunopathology and autoimmunity.

## Methods

### Mouse Models

All animal experiments were approved by the University Committee for Animal Resources at the University of Rochester. Only male wild-type (WT) and TNF-Tg (C57BL/6 background) mice were used for this study due to the accelerated mortality of females (Bell et al., 2019). The 3647-line of TNF-Tg mice (Douni et al., 1995-1996), originally obtained from Dr. George Kollias, have been maintained at the University of Rochester. Individual limbs were used as the experimental unit based on the asymmetry of TNF-Tg arthritis (Bell et al., 2019; Bouta et al., 2017; Kenney et al., 2022a; Li et al., 2013; Li et al., 2010; Li et al., 2011; Proulx et al., 2007). A total of 41 mice were used in this study, and quantitative metrics were evaluated with a minimum of n = 6 experimental units and n = 3 for qualitative histology, similar to previous studies (Bouta et al., 2017; Wu et al., 2019). Specific sample sizes for each experiment are provided in the Supplementary Methods. All mice were used for the study except three Advanced ankle joints were excluded from histologic assessment due to insufficient decalcification; otherwise, no mice or samples were excluded from the study. No specific inclusion or exclusion criteria were set. Animals were allocated into experimental groups based on age. Previously, we demonstrated that the joint disease in TNF-Tg mice can be defined as “Early” by mild synovitis and bone erosions associated with PLN expansion (increased volume and blood flow), and intact PLV contraction frequency and clearance in male mice at 5- to 6-months of age. On the other hand, “Advanced” joint disease occurs concomitant with PLN collapse (reduced volume and blood flow) and failure of PLV contractility that is commonly observed in male mice starting at 8-months of age (Bouta et al., 2018; Bouta et al., 2017; Li et al., 2013; Li et al., 2010; Li et al., 2011; Proulx et al., 2007; Scallan et al., 2021). All mice were housed in the same traditional caging environment and exposed to a 12-hour light/dark cycle to limit potential confounders. TNF-Tg mice were monitored weekly for failure to thrive and were provided diet supplementation with Nutra-Gel (2-ounce cups, Bio-Serv). There were no unexpected adverse events for the mice allocated to this study. Investigators performing histologic stains and histomorphometric analysis were blinded to the group allocation of the samples while conducting the experiment.

### Spatial Transcriptomics Tissue Optimization

PLNs from WT males (n=2 mice, 4 PLNs) were harvested, embedded in optimal cutting temperature (OCT) compound (Cat#4583; Sakura, Osaka, Japan), and frozen in liquid nitrogen. Six tissue sections of 10μm were placed within individual capture areas of a 10X Genomics Optimization slide and stored at -80C. The tissue sections were then fixed in cold methanol and stained with H&E according to the Visium Methanol Fixation and H&E Staining guide (10X Genomics). An Olympus VS120 slide scanner was used for brightfield imaging of the H&E-stained sections. To evaluate the optimal permeabilization time, each capture area was permeabilized for a different time period (3, 6, 12, 18, 24, or 30 minutes) followed by reverse transcription with fluorescent nucleotides to label cDNA. Tissue sections were then removed, and the slide was imaged again using the Olympus VS120 slide scanner to evaluate fluorescence with the TRITC filter. The optimal permeabilization time was determined to be 12-minutes by balancing the brightest fluorescent signal intensity with the least amount of diffusion (Supplementary Figure 1), and the 12-minute permeabilization time was then applied to the sections of the gene expression slide.

### Spatial Transcriptomics Gene Expression

PLNs from 5-6-month-old WT (n=5 mice, 10 PLNs), 5-6-month-old TNF-Tg (Early arthritis; n=3 mice, 6 PLNs), and >8-month-old TNF-Tg (Advanced arthritis; n=6 mice, 12 PLNs, 2 replicate tissue blocks of 6 PLNs each) mice were harvested, embedded in OCT, and frozen in liquid nitrogen. The PLNs were patterned in the block for post-hoc animal and limb identification. To evaluate RNA integrity numbers (RIN) for quality control, ten-consecutive 10μm sections were collected with a minimum RIN value of 9.6 for the gene expression samples (Supplementary Table 1). A 10μm tissue section was then placed within the 4 capture areas of a 10X Genomics Gene Expression slide and stored at -80C. The tissue was then fixed with cold methanol and H&E stained based on the Visium Methanol Fixation and H&E Staining guide (10X Genomics). An Olympus VS120 slide scanner was used for brightfield imaging of the slide at 20X, and .tif files were saved for downstream analysis in the SpaceRanger software. The tissue sections were then permeabilized for 12-minutes to release poly-adenylated mRNAs, which were captured by primers on the slide. The Visium Spatial Gene Expression Kit was then used for reverse transcription to produce full-length cDNA. The cDNA was assessed by KAPA qPCR to optimize the cycle numbers for cDNA amplification. Following amplification, the cDNA was purified using SPRISelect beads (Agilent, Santa Clara, CA). The Visium Library Construction Kit (10X Genomics) was then used to generate Illumina-compatible sequencing libraries with approximately 25% of the total cDNA yield. cDNA amplicon size was optimized, and indexed sequencing libraries were constructed by End Repair, A-tailing, Adaptor Ligation, and PCR. The NovaSeq 6000 sequencer (Illumina, San Diego, CA) was used to sequence the amplified libraries on an SP flow cell of 28×10×10×90 to obtain approximately 50,000 reads per spot for each capture area covered by tissue, according to the sequencing requirements in the Visium Spatial Gene Expression manual.

### Spatial Transcriptomics Analysis

The filtered feature matrices were imported and analyzed using the Seurat packages (v4.0.3) in RStudio (v1.2.1335; R v4.1.1) (Hao et al., 2021). The data was then processed according to instructions of the Seurat R package, where a shared nearest neighbor (SNN) clustering algorithm was used to embed the data as a Uniform Manifold Approximation and Projection (UMAP). Spot clusters were defined using resolution = 0.5, and both unsupervised and minimally supervised clustering (unsupervised plus manual combination of related clusters) was performed to group related spots based on their spatial location in the PLN. The different capture areas were merged to define relative spot counts of related clusters. Both capture areas for the PLNs from mice with Advanced arthritis were grouped together for analysis based on comparable spot expression via integration and clustering (Supplementary Figure 2). For analysis of individual PLNs, Loupe Browser (v5.0.1, 10X Genomics) was used to manually select PLN-specific spots. The selected PLN spots were exported as separate csv files and imported into Seurat for downstream analysis. The *AverageExpression* function in Seurat was also used to quantify the expression of particular genes within specific spatial clusters in the individual PLNs. Note that the spatial transcriptomics dots are not at single-cell resolution, but instead represent 55μm diameter spots for spatially-mapped bulk-sequencing of all cells within a particular spot. Thus, to estimate IgG expression by single B-cells within the sinuses by spatial transcriptomics, we normalized to *Ighm*, which is highly expressed by all B-cells via scRNAseq (Figure 5).

### Single Cell Sequencing and Analysis

PLNs from TNF-Tg mice with Early and Advanced arthritis (n = 3 mice pooled per group) were harvested and surrounding tissue was removed. The PLNs were incubated in 10% fetal bovine serum (FBS) in Dulbecco’s modified Eagle’s medium (DMEM) + GlutaMAX (Gibco, Cat# 10566-016), followed by mechanical disruption of tissue placed in a petri dish with 1.5mL of Accumax (Innovative Cell Technologies, Cat# AM-105) with a scalpel. Minced tissue in Accumax was then transferred into a 1.5mL Eppendorf tube and rotated for 1-hour at room-temperature. After the digestion, single cells were passed through a 70μm MACS SmartStrainer (Miltenyi Biotec, Cat# 130-098-462) that was pre-wet with 3mL of 20% FBS in PBS into a 15mL conical tube on ice. 1mL of 20% FBS was used to clean the Eppendorf tube then similarly passed through the cell strainer to quench the enzymatic reaction. All samples for a particular group were pooled together, pelleted at 300g and incubated at 4°C for 10-min, and resuspended in 2% FBS in PBS. The cell suspension was stained with Hoechst 33342 (exclude red blood cells; NucBlue Live ReadyProbes, ThermoFisher Scientific, Cat# R37605) and Sytox Green (exclude dead cells; NucGreen Dead 488 ReadyProbes, ThermoFisher Scientific, Cat# R37109). An aliquot was set aside as an unstained control. Live cells were then sorted on a BD FACSAria II using a 100μm nozzle into a 2mL Eppendorf collection tube that had been filled completely overnight with 100% FBS, and replaced with 200μL of fresh 2% FBS collection media prior to sorting. A total of 100,000 Hoechst^+^/Sytox^−^ events were collected for each group and processed for single-cell RNA-sequencing by the Genomics Research Center at the University of Rochester Medical Center. The cells were sequenced using Illumina’s NovaSeq 6000.

The datasets were analyzed according to the instructions of the Seurat package in R with unsupervised SNN clustering performed at resolution = 0.5. For the analysis, features were removed if the gene was only expressed by < 3 cells, and low-quality cells were removed if the cell exhibited < 200 features. Differential gene expression analysis was evaluated and used for annotation of each cluster. To determine potential cell interactions, the NicheNet (v1.0.0) (Browaeys et al., 2019) R package was applied to the Seurat objects using commands available through SeuratWrappers (v0.3.0), and only *bona fide* interaction potential was evaluated (i.e., supported by literature). A published dataset of LEC single-cell RNA-sequencing ((Xiang et al., 2020), GSE145121) was imported into R and integrated with our spatial transcriptomics to assess specific LEC populations within our spatial dataset. The published data were reclustered to identify the LEC subpopulations. Then, overlap of gene expression between the spatial spots (multiple cells) and the single-cell data generated a prediction score of cell localization to specific spatial spots. Analysis was performed according to the instructions of the “Analysis, visualization, and integration of spatial datasets with Seurat” vignette provided by the Satija laboratory.

### Micro-CT Data Collection and Analysis

*Ex-vivo* micro-CT on the hind paws was performed to evaluate and correlate the severity of ankle arthritis with gene expression in the efferent PLN. Micro-CT datasets were acquired using a VivaCT 40 (Scanco Medical, Bassersdorf, Switzerland) with the following imaging parameters: 55kV, 145μA, 300ms integration time, 2048 × 2048 pixels, 1000 projections over 180°, resolution 17.5μm isotropic voxels. The DICOM files were then imported into Amira software (v2020.2, ThermoFisher Scientific, FEI, Hillsboro, OR, USA) and talus bone volumes were measured, as previously described (Kenney et al., 2022b; Proulx et al., 2007).

### Histology, Immunostaining, and Image Analysis

PLNs from additional Early and Advanced TNF-Tg mice (n = 3 per group) were harvested for the paraffin-embedded histology, while the ankle joints from the mice used for spatial transcriptomics were utilized. Due to inefficient decalcification and deficient soft-tissue removal, ankle joints from n = 3 Advanced mice were excluded, and the ankle joints from the mouse with Advanced arthritis used for PLN 3D modeling was used for histology. PLNs and ankle joints were fixed in 10% neutral buffered formalin (NBF) for 2h and 3-days, respectively. PLNs were then embedded in paraffin, while ankles were decalcified in Webb-Jee 14% EDTA solution for 1-week prior to paraffin-embedding. Tissue sections were then collected at 5μm. The human synovial samples were processed according to the Accelerating Medicines Partnership (AMP) Program’s standard operating procedures (Der et al., 2019; Lewis et al., 2019; Wei et al., 2020; Zhang et al., 2021; Zhang et al., 2019).

For Prussian Blue staining, both frozen sections from the spatial transcriptomics tissue blocks and paraffin-embedded sections were evaluated. The frozen PLN spatial transcriptomics sections were used for quantification of Ferritin^+^ cells, while the PLN paraffin-embedded sections were used for representative images of the Prussian blue stain. Frozen sections were first fixed in chilled 10% NBF for 10-minutes then stained with Prussian Blue and counterstained with Nuclear Fast Red. The paraffin-embedded sections were dewaxed and rehydrated before the staining process. Paraffin-embedded ankle sections were also stained with Prussian Blue. Each slide was then imaged using a VS120 Slide Scanner, and .vsi files were analyzed using semi-automated segmentation workflows in Visiopharm (v2021.07; Horsholm, Denmark).

For immunofluorescence staining, slides were incubated at 60°C overnight to melt the paraffin and hydrated by immersion in xylenes, graded alcohols and water. Antigen retrieval was performed by boiling slides in a Coplin jar filled with Citrate-Based Antigen Unmasking Solution (1:100 dilution; Vector Laboratories, Cat# H-3300). After cooling to room temperature, non-specific staining was prevented by incubating tissues with normal donkey serum (NDS; Jackson ImmunoResearch, Cat# 017-000-121) in TBS (Bio-Rad, Cat# 1706435) / 0.3% Triton X-100 (Millipore Sigma, Cat# X100) for 40-min at room temperature. The primary antibodies were then diluted in the NDS solution and applied at 4°C overnight. After three PBS washes, secondary antibodies were similarly diluted in the NDS solution and incubated on the slides at room temperature for 1-hour. The slides were then washed with PBS three times and mounted using one drop of NucBlue Live ReadyProbes (Hoechst 33342) and ProLong Gold Antifade Mountant (ThermoFisher Scientific, Cat# P36930). Finally, the slides were imaged using a VS120 Slide Scanner or a Zeiss Axioplan Microscope.

Antibodies and dilutions for immunofluorescent staining: Rabbit anti-mouse CD6 (ThermoFisher, Cat# MA5-29680, RRID: AB_2785505, 1:50), rat anti-mouse F4/80 (BioRad, Cat# MCA497R, RRID:AB_323279, 1:50), rabbit anti-mouse Marco (Abcam, Cat# ab239369, 1:100), Alexa Fluor 555 goat anti-mouse IgG2b cross-adsorbed (ThermoFisher Scientific, Cat# A-21147, RRID:AB_2535783, 1:50), APC rat anti-mouse CD45R/B220 (BD Biosciences, Cat# 553092, RRID:AB_398531, 1:100), goat anti-mouse Alcam (CD166; ThermoFisher, Cat# PA5-47083, RRID:AB_2607383, 1:100), Cy3 goat anti-mouse IgM (Jackson ImmunoResearch Laboratories, Cat# 115-165-020, RRID:AB_2338683, 1:200), FITC donkey anti-mouse IgG (Jackson ImmunoResearch Laboratories, Cat# 715-095-150, RRID:AB_2340792, 1:200), goat anti-proliferating cell nuclear antigen (PCNA; Santa Cruz Biotechnology, Cat# sc-9857, RRID:AB_2160372, 1:50), FITC Peanut Agglutinin (PNA; Millipore Sigma, L7381-1MG, 1:50), goat anti-mouse peripheral node addressin (PNAd; BD Biosciences, Cat#553863, RRID:AB_395099, 1:50), rabbit anti-mouse Lyve1 (Acris Antibodies, Cat# DP3513, RRID:AB_1004776, 1:50), AlexaFluor 568 donkey anti-goat IgG (ThermoFisher Scientific, Cat# A-11057, RRID:AB_142581, 1:200), AlexaFluor 647 F(ab’)_2_ fragment donkey anti-rat IgG (Jackson ImmunoResearch, Cat# 712-606-153, RRID:AB_2340696, 1:200), and AlexaFluor 647 donkey anti-rabbit IgG (Abcam, Cat# ab150075, RRID:AB_2752244, 1:400).

### Human Synovial Samples

Human synovial samples were provided by JHA from arthroplasty or synovectomy tissues (RSRB00055411) collected using Accelerating Medicines Partnership (AMP) standard operating procedures (Der et al., 2019; Lewis et al., 2019; Wei et al., 2020; Zhang et al., 2021; Zhang et al., 2019). Prussian blue staining was evaluated in synovial samples from patients with osteoarthritis (OA, n = 3) and RA (n = 3) (fulfilling 2010 ACR/EULAR classification criteria). OA samples had limited Prussian Blue staining and the region containing the maximum iron-laden cells is depicted in Figure 4. All of the RA samples were derived from CCP+ subjects. Two RA samples looked similar to OA, with limited to no iron-laden cells. One of these RA samples did not have a histology pathotype (Lewis et al., 2019) assigned and lacked information about treatments. The second RA sample had a lymphoid pathotype, and the patient received infliximab. The RA sample depicted in Figure 4 exhibited regions of abundant iron-laden cells. This synovectomy tissue was of lymphoid pathotype obtained from a subject treated with prednisone and methotrexate, but with persistently high DAS28-CRP of 4.9.

### PLN 3D Modeling

A WT, Early, and Advanced PLN (n=1/group) was sectioned at 5μm through the entire lymph node. Each section was then stained with Prussian Blue and Nuclear Fast Red counterstain, then imaged using a VS120 slide scanner. Images (.tif) were exported from the slide scanner, and image stacks were imported into Amira software. The “Align Slices” module was used to rotate and align each individual section. The images were also imported into ImageJ for color segmentation using the “Color Deconvolution 2” plug-in, where blue (iron) and red (stromal cells) were segmented using the Feulgen LightGreen vector with a set threshold. The segmented red and blue sections were then exported as an image sequence, and then imported into Amira and aligned with the original images. Animations were then generated in Amira to develop the Supplementary Videos.

### Flow Cytometry

Single-cell suspensions were collected from PLNs, spleen, whole blood, and bone marrow of TNF-Tg mice with Early and Advanced arthritis (n = 3 mice per group). A lethal dose of ketamine / xylazine cocktail was administered intraperitoneal prior to tissue harvest. Evan’s Blue dye (2%) was injected intradermal into the hindpaws of the mice for identification of the PLNs, then the fur on the posterior aspect of the hindlimb was removed with depilatory cream. The blood was collected by cardiac puncture and placed into a 1.5mL Eppendorf tube with 5mM EDTA (pH 8.0, RNase-free; ThermoFisher Scientific, Cat# AM9261). For the PLNs and spleen, the organs were incubated in 10% FBS in DMEM prior to removal of external tissue and single-cell isolation by mechanical digestion of the tissue incubated in 3mL of 20% FBS in PBS using a scalpel. The bone marrow was then collected from the femurs, as previously described (Amend et al., 2016). Briefly, the femurs were dislocated with the condyles and epiphyses removed. An 18-gauge needle was used to puncture a hole through a 0.65mL Eppendorf tube, which was then placed inside a 1.5mL Eppendorf tube. The femurs were then placed inside the 0.65mL Eppendorf tube, centrifuged at 10,000g for 15-sec, and a pellet was formed in the 1.5mL Eppendorf tube. The bone marrow pellet was then resuspended in 20% FBS.

For all organs, a 30μm MACS SmartStrainer (Miltenyi Biotec, Cat# 130-098-458) was pre-wet was 3 mL of 20% FBS. Next, the cell suspensions were passed through and 20% FBS was added until the total volume reached 10mL. Each sample was then pelleted at 300g and 4°C for 10min and resuspended in red blood cell (RBC) lysis buffer (ThermoFisher Scientific, Cat# 00-4333-57). The PLNs and bone marrow were incubated in 5mL of lysis buffer, while the spleen and blood were resuspended in 10mL of lysis buffer for 5min at room temperature. The lysis was then stopped by adding PBS up to 40mL to each sample. The cells were then pelleted at 300g and 4°C and resuspended in staining buffer (1% bovine serum albumin (BSA) in PBS). Data from 2-million cells per sample was collected using an Aurora Full Spectrum Flow Cytometer (Cytek Biosciences) and analyzed using FlowJo (v10.8.1; BD Biosciences). Prior to the flow cytometry experiments, optimal antibody titrations and compensation matrix were setup using single-stain controls on cells isolated from the PLNs and spleen of an Advanced TNF-Tg mouse (n=1) using the protocols described above.

For the staining, the antibodies were diluted in brilliant stain buffer (BD Biosciences, Cat# 563794, 1:4) / 1% BSA staining buffer. The following antibodies and dilutions were used: AlexaFluor 488 CD45R/B220 (Biolegend, Cat# 103225, 1:160), AlexaFluor 700 CD20 (Biolegend, Cat# 150416, 1:40), AlexaFluor 647 CD19 (Biolegend, Cat# 115522, 1:160), BrilliantViolet 785 CD3 (Fisher Scientific [Biolegend], Cat# 50-402-911, 1:20), BrilliantViolet 711 CD11b (Biolegend, Cat# 101242, 1:80), BrilliantViolet 650 IgM (Fisher Scientific [BD Biosciences], Cat# BDB564027, 1:20), BrilliantViolet 605 IgG (Biolegend, Cat# 405327, 1:40), BrilliantViolet 421 CD138 (Biolegend, Cat# 142508, 1:80), PE-Cy7 CD23 (Biolegend, Cat# 101614, 1:160), 7-AAD (ThermoFisher, Cat# 00-6993-50, 1:20), PE/Dazzle 594 CD21 (Biolegend, Cat# 123440, 1:160), and AlexaFluor 555 IgG2b (ThermoFisher, Cat# A-21147, 1:40).

### IgG2b ELISPOT

PLNs were harvested from TNF-Tg mice with Early and Advanced arthritis (n=3 mice, 6 PLNs per group). Single-cell suspensions and red blood cell lysis was performed as in the Flow Cytometry methods. The isolated cells were then processed using the directions of the Mouse IgG2b Single-Color ELISPOT kit without *in vitro* stimulation (Immunospot). The wells were imaged using an ELISPOT reader (Immunospot), and spots were counted manually after importing the images into ImageJ.

### IgG Isotyping Serum ELISA

Blood from the Early and Advanced TNF-Tg mice used in the scRNAseq (n = 3 mice / group) was collected by cardiac puncture. The blood was allowed to sit at room temperature for 30-min and then was centrifuged at 1500g for 10-min at 4°C. The serum was then collected, and aliquots were frozen at -80°C. Serum was then processed and analyzed using the instructions of the Mouse IgG Isotyping kit (Abcam, Cat# ab273149).

### Statistical Analysis

Linear regression, t-test, one-way ANOVA, and two-way ANOVA statistical analyses were performed in GraphPad Prism (v9.1.0, San Diego, CA, USA).

## Results

### Spatial Transcriptomics Identifies PLN Sinus Regions with Enhanced Immunoglobulin Production in TNF-Tg Mice with Advanced Arthritis

As dynamic cellular changes have been identified in the PLN sinuses of TNF-Tg mice (Li et al., 2011), we utilized a spatial transcriptomics approach to specifically target gene expression changes in these histologically defined sinus regions. To facilitate visualization of spatial gene expression, representative H&E-stained images of the PLN tissues and clustering of spatial transcriptomics datasets from wild-type and TNF-Tg mice with Early and Advanced arthritis are presented in Figure 1A-D. Uniform manifold approximation and projection (UMAP) plot of the merged datasets with minimally supervised clustering (resolution = 0.5 and manual combination of clusters with similar anatomical localization) reveals 3 predominant anatomical regions of the PLNs: paracortex, follicles, and sinuses, based on their gene expression profiles and spatial location (Fig. 1E). Note the close association of the sinus spots (red) mapped to the regions of the PLN with reduced cell density (white areas of H&E images) representing the PLN sinuses (Figs. 1A-D). The spatially localized gene expression analysis identified the PLN sinuses as *Lyve1*^Hi^/*Marco*^Hi^ (Figs. 1F,G), consistent with markers of lymphatic endothelial cells (LECs) described in prior transcriptomic studies (Xiang et al., 2020). The cell-dense regions of the paracortex were populated by hematopoietic cells based on high expression of *Ptprc* (CD45)^Hi^ (Fig. 1H), and the B-cell follicles were enriched with *Ptprc*^Hi^/*Ms4a1* (CD20)^Hi^ expression (Fig. 1I). Further unsupervised clustering (resolution = 0.5) identified 11 unique spot clusters where the sinus regions of the PLN (within blue-dashed line) contained two predominant clusters: Sinus-Immuno (immunoglobulin enriched) and Sinus-LEC (Fig. 1J). The Sinus-Immuno cluster demonstrated enrichment of immunoglobulin gene expression, such as *Ighg1, Ighg2b, Ighg2c*, and *Ighm* (Fig. 1K-N). In contrast, the Sinus-LEC cluster was defined by relatively increased expression of *Lyve1* (Fig. 1F) and *Marco* (Fig. 1G). These predominant sinus clusters were then subset for each condition (Fig. 1O) to quantify and compare the spot abundance of Sinus-Immuno vs Sinus-LEC. PLNs from both TNF-Tg mice with Early and Advanced arthritis exhibited an increased number of sinus spatial spots compared to WT (Fig. 1P; WT 20.0±8.0, Early 56.2±37.6, Advanced 77.3±32.0 sinus spots), while Advanced PLNs showed a greater proportion of Sinus-Immuno spots compared to Early PLNs (Fig. 1Q; WT 64.6±7.8, Early 43.1±27.0, Advanced 71.6±17.7% Sinus-Immuno spots).

**Figure 1.**
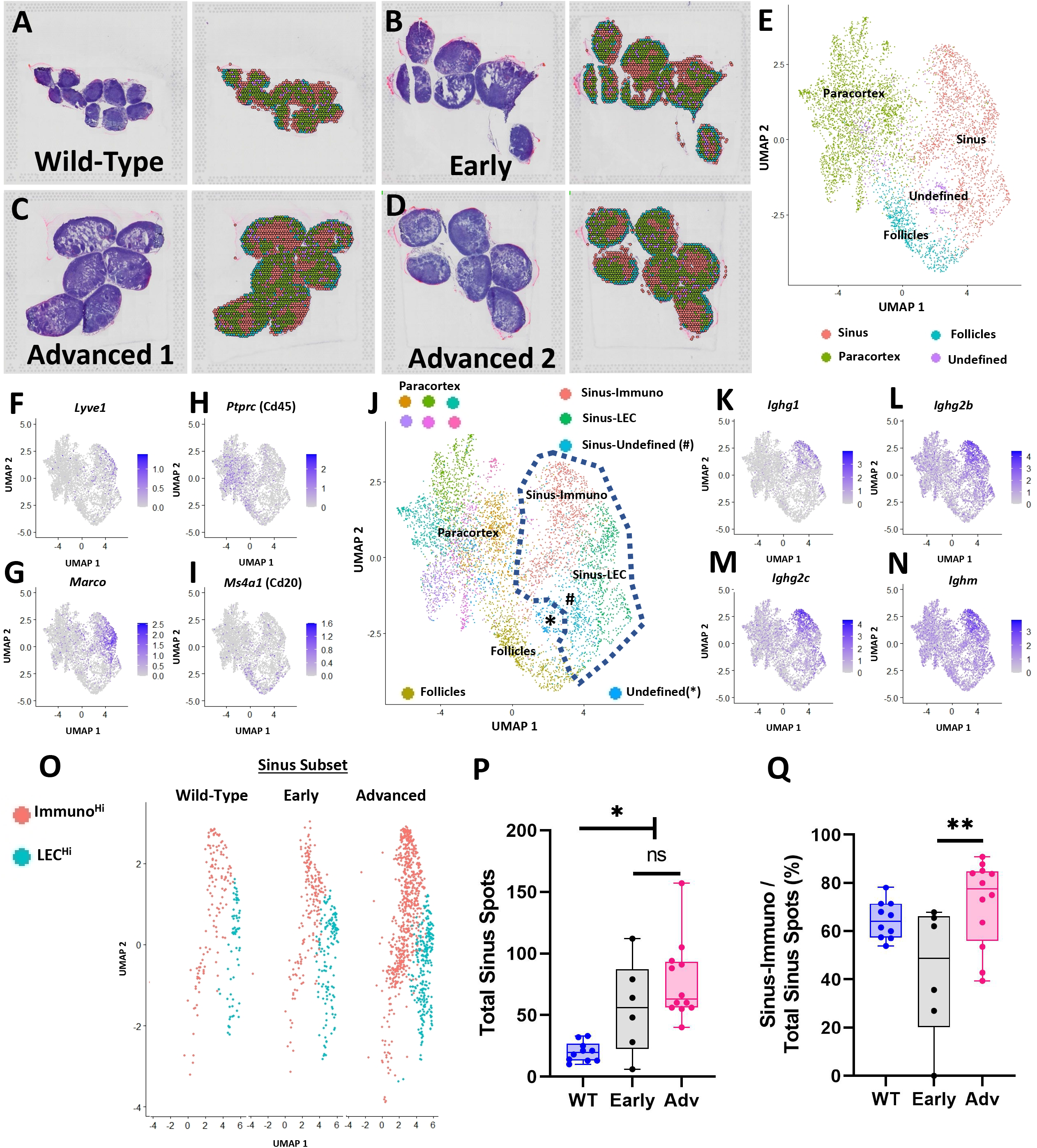
Spatial transcriptomics identifies PLN sinus regions with enhanced immunoglobulin production in TNF-Tg Mice with Advanced arthritis. PLNs from WT (5.5-month-old; n=10 PLNs) **(A)** and TNF-Tg mice with Early (5-6-month-old; n=6 PLNs) **(B)** and Advanced (>8-month-old; n=12 PLNs, 6/capture area) **(C-D)** arthritis were harvested and processed for spatial transcriptomics. Each H&E-stained image (left) corresponds to a transcriptional representation (right) of the PLN sinus (red spots overlaying white sinus spaces), paracortical (green spots), and follicular (blue spots) regions. Isolated spots with high ribosomal gene content were annotated as an undefined (purple) cell population **(A-E)**. Feature plots represent the identification of *Lyve*^Hi^*/Marco*^Hi^ sinuses **(F-G)**, *Ptprc*^Hi^ paracortices **(H)**, and *Ptprc*^Hi^*/Ms4a1*^Hi^ follicles **(I)** that were validated by histologic localization as in **A-D**. Unsupervised clustering (resolution=0.5) resolved 11 transcriptionally distinct PLN regions, where the sinus-associated spots localized to 3 distinct clusters (dashed lines) **(J)**. The *Ighg1*^Hi^/*Ighg2b*^Hi^/*Ighg2c*^Hi^/*Ighm*^Hi^ Sinus-Immuno **(K-N)** and *Lyve1*^Hi^/*Marco*^Hi^ Sinus-LEC **(F-G)** populations predominated, while Sinus-Undefined spots exhibited a high proportion of ribosomal genes. The predominant Sinus-Immuno (red) and Sinus-LEC (blue) populations were subset and re-clustered **(O)**, and individual PLNs were evaluated for total sinus-associated spots (Sinus-Immuno + Sinus-LEC) **(P)** and proportion of Sinus-Immuno spots **(Q)**. Statistics: One-way ANOVA, *\*p<0*.*05, **p<0*.*01* **(P**,**Q)**.

### PLN Expansion of Marco^+^ Peri-Follicular Medullary Sinuses Determined by Spatial and Single-Cell Integration

Due to the dynamic cellular changes within the TNF-Tg PLN sinuses during the onset and progression of inflammatory-erosive arthritis, we aimed to elucidate the identity of the affected sinus regions. To localize the expanded TNF-Tg PLN sinus regions with distinct LEC populations, we integrated our spatial transcriptomics with a published scRNAseq dataset of lymph node LEC subtypes (Xiang et al., 2020) (Fig. 2A), including those localized to the *Ackr4*^+^ subcapsular ceiling (cLECs), *Madcam1*^+^ subcapsular floor (fLECs), *Ccl21a*^*+*^ collecting vessel, *Marco*^*+*^ peri-follicular medullary, and *Ptx3*^*+*^ central medullary sinuses (Fig. 2B-F). We then computed prediction scores for each spatial spot corresponding to the LEC subpopulations generated from scRNAseq dataset by using *FindTransferAnchors* function in Seurat and visualized each LEC subtype on the UMAP defined by the spatial transcriptomics analysis (Fig. 2G-J). The Marco-LECs were associated with the expanded sinus regions (Fig. 2K, red spots within dashed lines). This spatial identity of increased Marco^+^ sinuses in TNF-Tg lymph nodes was confirmed by histology with sinus regions (Fig. 2L,M, red spots; reproduced from Fig. 1D for convenience) that showed enrichment for Marco-LECs (Fig. 2N, yellow/red expression) relative to other LEC subtypes (Fig. 2O-Q). Quantitative analysis confirmed a significant enrichment for Marco-LECs within the spatially-defined sinus regions (Fig. 2R) but not in paracortex and follicle regions (Fig. 2S-U). Thus, by integrating scRNAseq and spatial transcriptomics datasets, we identified the Marco^+^ peri-follicular medullary sinuses as those involved in PLN “Expansion” and “Collapse” during TNF-Tg arthritis progression.

**Figure 2.**
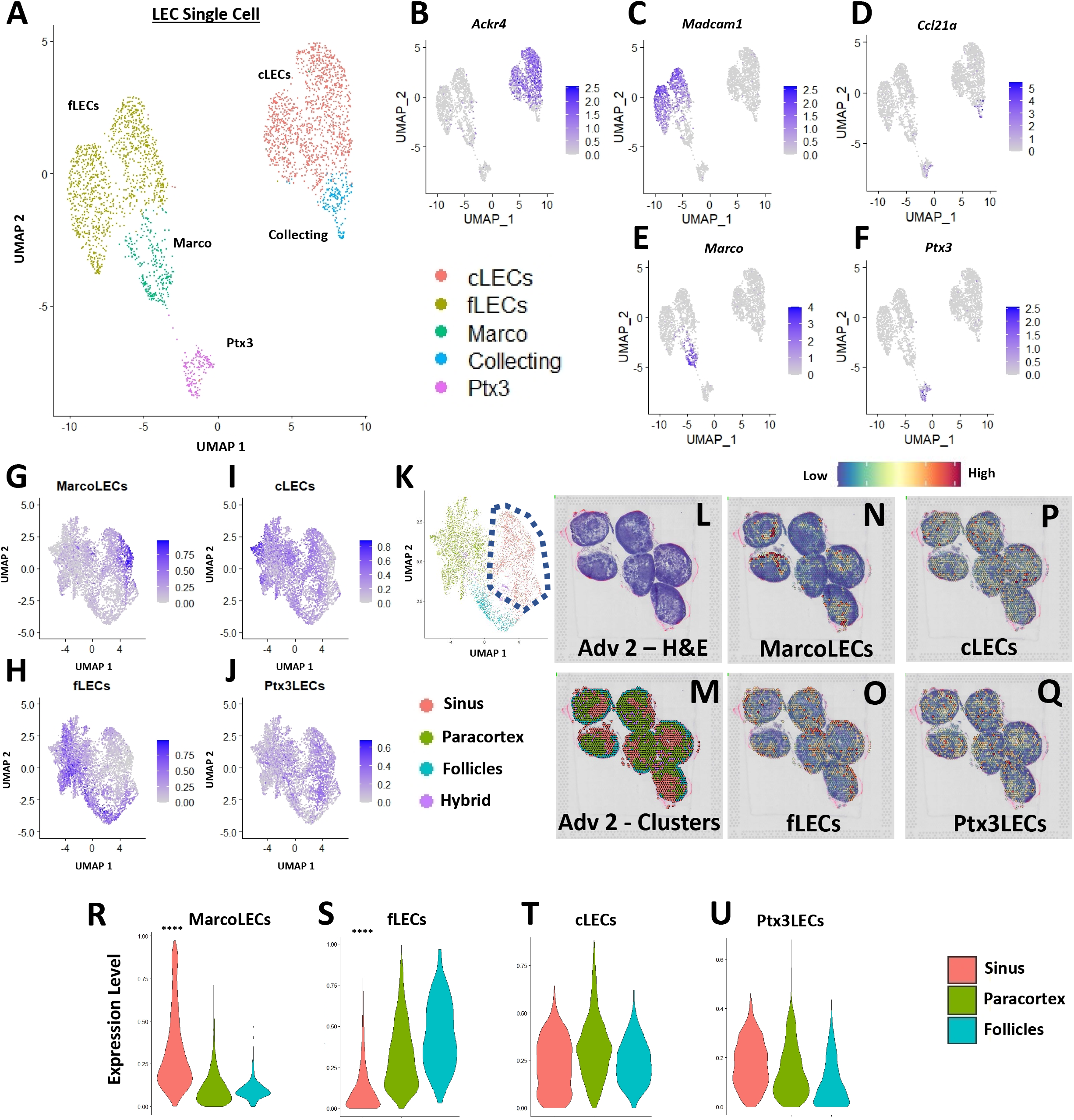
*Marco*-expressing peri-follicular medullary LECs predominate in the spatially defined sinus regions. Subpopulations of LECs from a published scRNAseq dataset of C57BL/6 murine lymph nodes ((Xiang et al., 2020), GSE145121) were visualized in a UMAP **(A)**. Specific LEC populations were defined by gene enrichment **(B-F)**, as previously described (Xiang et al., 2020). The LEC single-cell data was integrated with the regions identified by spatial transcriptomics, where feature plots demonstrate the selective enrichment for Marco-LECs in the sinuses **(G)**, fLECs in the paracortices and follicles **(H)**, and the general distribution of cLECs **(I)** and Ptx3-LECs **(J)** throughout the spatially-resolved PLN regions **(K)**, identified in Figure 1. A representative H&E-stained image of the Adv 2 capture area **(L)** with cluster annotation **(M)** is shown as a spatial feature plot to further demonstrate the Marco-LECs localized to the expanded sinus regions of TNF-Tg PLNs **(N-Q)**. Quantification of Marco-LECs **(R)**, fLECs **(S)**, cLECs **(T)**, and Ptx3-LECs **(U)** within the spatially-resolved PLN regions confirmed a significant enrichment for Marco-LECs (Log2FC = 0.354) within and fLECs (Log2FC = -0.321) outside of the PLN sinuses. Statistics: Wilcoxon Rank-Sum Test of sinus vs. all other regions; *\*\*\*\*FDR < 2*.*23E-308* **(R, S)**.

### Aggregation of IgG2b^+^ Plasma Cells Adjacent to PLN Marco^+^ Sinuses Correlates with Reduced Bone Volumes in the Afferent Ankle Joint

To examine gene expression changes within the Marco^+^ sinus regions of the TNF-Tg PLNs and severity of inflammatory-erosive arthritis in the afferent ankle joint, *ex-vivo* micro-CT of the hindpaws was performed (Fig 3A-B; note the dislocation of talus bone from tibia in Advanced arthritis). As expected, the mice with Advanced arthritis had significantly decreased talus bone volumes compared to mice with Early arthritis (Fig. 3C; Advanced 0.40±0.15 vs Early 0.90±0.20mm^3^, *p<0*.*0001*). The spatial transcriptomic spot-annotated images of the associated PLNs from TNF-Tg mice with Early (Fig. 3D) and Advanced (Fig. 3E, Advanced 2 capture area as representative) arthritis are shown to identify the sinus regions (red spots; reproduced from Fig. 1B,D for convenience). We next examined spatial expression of *Ighg2b* (Figs. 3F,G; blue = low; white = medium; red = high expression) in PLNs with notable expression of *Ighg2b* within the sinus regions. Furthermore, PLN sinuses from TNF-Tg mice with Advanced arthritis showed significantly greater *Ighg2b*/*Ighm* expression ratio (see Supplementary Methods, Spatial Transcriptomic Analysis) compared to those with Early arthritis (Figs. 3H; Early 0.52±0.12 vs Advanced 1.41±0.45 counts/counts, *p<0*.*001*). Importantly, expression levels of *Ighg2b* demonstrated a significant negative correlation with talus bone volumes (Fig. 3I; R^2^=0.54, *p=0*.*0005*), suggesting a strong relationship between arthritic severity and immune responses in the joint-draining PLNs. *Ighg2b* interestingly exhibited a selective relationship with arthritis severity compared to other immunoglobulin genes, such as *Ighg1* and *Ighg2c* relative to *Ighm* (Supplementary Figure 3). Immunofluorescence showed a limited accumulation of IgG2b^+^ cells in Early TNF-Tg PLNs (Fig. 3J), while IgG2b^+^/B220^−^ cells located in the Marco^+^ sinus tissue significantly increased in Advanced PLNs (Fig. 3K), confirming the results of the spatial transcriptomics analysis (Fig. 3L; Early 15.9±2.3 vs Advanced 40.3±8.1 IgG2b^+^ cells / 200x field, *p<0*.*0001*). IgM^+^ cells were also found to be significantly reduced in Advanced PLNs, associated with the increase in IgG^+^ cells and reduced cell proliferation by PCNA staining, supporting plasma cell identity (Supplementary Figure 4). Flow cytometry confirmed a significant increase in IgG2b^+^/CD138^+^ plasma cells in Advanced PLNs (Figs. 3M-O; Early 11.4±3.1% vs Advanced 55.6±4.6% of IgG2b^+^/CD138^+^ relative to CD11b^−^/CD3^−^ live cells, *p<0*.*001*; gating shown in Supplementary Figure 5). Increased IgG2b^+^ cells were also found systemically (bone marrow, blood, and spleen) of Advanced mice, although the significant increase in CD138^+^ plasma cells was selective for PLNs (Supplementary Figure 6 and Supplementary Table 2). Advanced PLNs also exhibited an increase in IgG2b secreting cells by ELISPOT (Figs. 3P-R; Early 55.8±14.6 vs Advanced 225.8±135.4 spot-forming units per million cells, *p<0*.*05*). Remarkably, despite an increase in plasma and IgG2b-secreting cells, there was no difference in IgG2b antibody levels in the serum of Advanced mice measured by ELISA (Supplemental Figure 6). The IgG2b isotype was also the predominant immunoglobulin in both Early and Advanced mice, as previously reported in C57BL/6 mice (Klein-Schneegans et al., 1989). Together, these data suggest that plasma cell differentiation and IgG2b class-switching in joint-draining lymph nodes are integral to the onset of Advanced arthritis in TNF-Tg mice.

**Figure 3.**
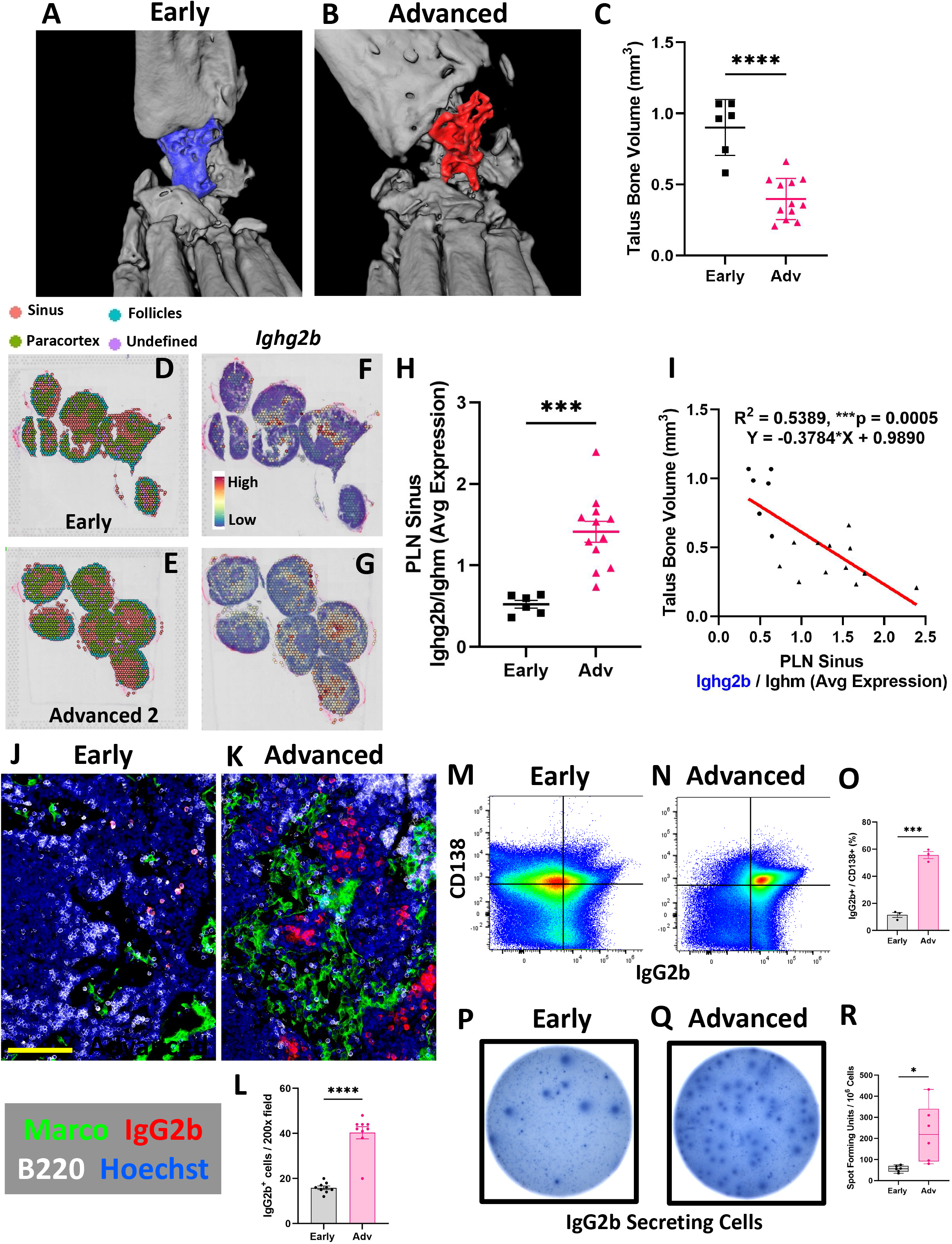
Increased IgG2b expression in PLN sinuses correlates with reduced bone volumes in the afferent ankle joint. *Ex-vivo* micro-CT of TNF-Tg hindpaws with Early **(A)** and Advanced **(B)** arthritis showed significantly reduced talus bone volumes in Advanced joints **(C)**. A transcriptional representation of the sinus regions (red) for Early **(D)** and Advanced (i.e. Adv 2) **(E)** PLNs is shown with *Ighg2b* expression depicted as a spatial feature plot (**F**,**G**; red = high, blue = low expression). There was a significant increase in *Ighg2b/Ighm* expression ratio in Advanced vs Early PLNs **(H)**, along with a significant and negative correlation of *Ighg2b* sinus expression with talus bone volumes (R^2^ = 0.54, *p<0*.*001*; circles = Early, triangles = Adv) **(I)**. The increase in IgG2b^+^ cells in Advanced TNF-Tg PLNs was validated by immunofluorescence where IgG2b^+^ cells are limited in Early PLNs **(J)**, while Advanced PLNs **(K)** exhibit a significant accumulation of IgG2b^+^ (red)/B220^−^ (white) plasma cells **(L)** localized to the Marco^+^ (green) PLN sinuses. Flow cytometry **(M-O)** and ELISPOT **(P-R)** confirmed significantly increased IgG2b expressing and secreting cells in Advanced PLNs, respectively. Statistics: Unpaired t-test **(C, H, L, O, R)**; linear regression **(I)**; \**p<0*.*05, ***p<0*.*001, ****p<0*.*0001*. Yellow scale bar = 100μm **(J**,**K)**.

### Accumulation of Synovial and PLN Sinus Ferric Iron-Laden Macrophages Associated with Decline in Tissue Fth1 Expression and Advanced Arthritis

To further elucidate the cellular dynamics within the TNF-Tg PLN sinus regions associated with the lymphocyte activation and plasma cell accumulation, we performed differential gene expression analysis between Early and Advanced cohorts. Specifically, ferritin heavy chain (*Fth1*) that encodes the heavy subunit of ferritin (the major iron storage protein) was significantly reduced in Advanced PLN sinuses relative to Early PLNs (Fig. 4A-C; Early 2.5±0.74 vs Advanced 1.0±0.50 counts, *p<0*.*001*). Additionally, *Fth1* expression levels were positively correlated with talus bone volumes in ipsilateral ankles (Fig. 4D, R^2^ = 0.47, *p<0*.*01*). To characterize the changes in Ferritin levels between Early and Advanced TNF-Tg PLNs, we performed Prussian blue with Nuclear Fast Red counterstain histochemistry. Compared to Early PLNs lacking Prussian blue^+^ cells (Fig. 4E,F), Advanced PLNs exhibited a significant accumulation of iron-laden cells localized to the sinuses (Fig. 4G-I, WT 0.043±0.014% vs Early 0.020±0.021% vs Advanced 0.37±0.36% Prussian blue to total PLN area, *p<0*.*05*). We also performed 3D reconstructions of a Prussian blue-stained PLN from each group and confirmed the global accumulation of iron-laden cells selective for Advanced PLNs (Supplementary Videos). As macrophages are critical for regulating iron homeostasis (DeRosa and Leftin, 2021), we further confirmed the identity of the Prussian blue^+^ cells as iron-laden macrophages by their immediate proximity to F4/80^+^/Marco^+^ macrophages aggregated within Marco-LEC lined PLN sinuses (Supplementary Figure 7). To evaluate the potential origins of these ferric iron (Fe^3+^) rich macrophages, we next assessed ankle joints by Prussian blue staining, which showed a dramatic increase in Fe^3+^-laden macrophages within the synovium of TNF-Tg mice with Advanced arthritis (Fig. 4J-N; WT 163.3±117.7 vs Early 7850±6090 vs Advanced 31897±16007 μm^2^ of Prussian blue in the peri-talus region; *p<0*.*001*). As a preliminary clinical correlate study of the Fe^3+^ macrophage phenotype, we assessed Prussian blue-stained human synovium. Relative to osteoarthritic synovium with a paucity of Fe^3+^ cells (Fig. 4O), ferric iron-laden macrophages were abundant within RA synovium during active disease (Fig. 4P, black arrow). Given the limited number of Fe^3+^-laden macrophages in Early arthritis in TNF-Tg mice, we conclude that inflammation alone is insufficient for the accumulation of ferric iron in macrophages, and hypothesize that additional inciting events (e.g., phagocytosis of dead and dying cells, initiation of ferroptotic processes) are necessary to promote their generation and accumulation.

**Figure 4.**
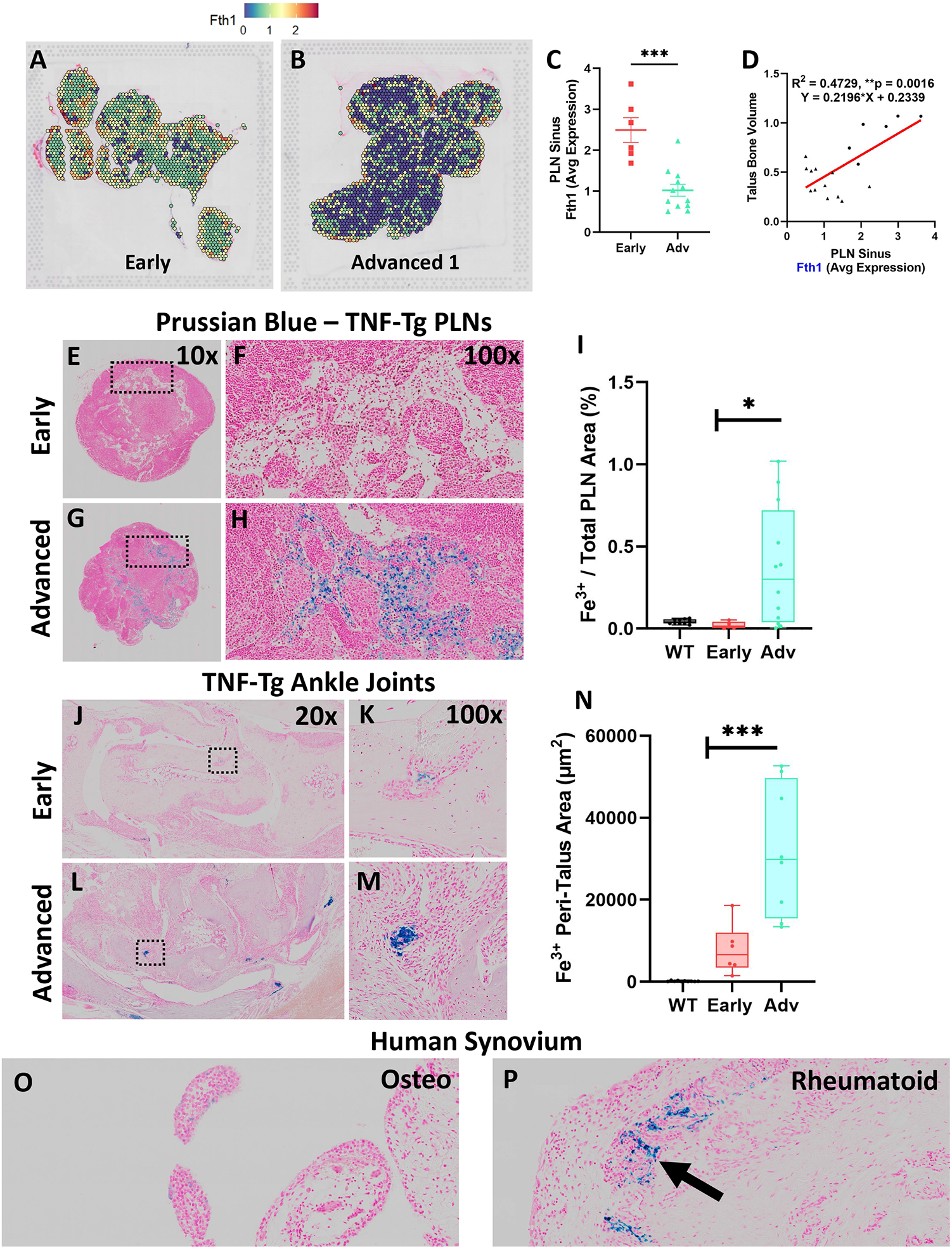
Accumulation of synovial and PLN sinus ferric iron-laden macrophages associated with decline in tissue *Fth1* expression and Advanced arthritis. Differential gene expression analysis within the PLN sinus regions revealed reduced expression of *Fth1* (ferritin heavy chain 1) in Advanced compared to Early PLNs (green to blue spots) **(A, B)**. Quantification of *Fth1* expression within the sinus regions of individual PLNs confirmed a significant reduction in Advanced PLNs **(C)**, and intra-sinus *Fth1* levels were positively correlated with talus bone volumes in the ipsilateral ankles (R^2^ = 0.47, *p<0*.*01*; circles = Early, triangles = Adv) **(D)**. Compared to Early PLNs with an absence of Prussian blue^+^ cells **(E, F)**, Fe^3+^-laden macrophages (blue) were significantly increased within the sinuses of Advanced PLNs (*p<0*.*05*) **(G-I)**. Similarly, the ankle synovium showed a significantly increased abundance of Fe^3+^ macrophages in Advanced versus Early TNF-Tg joints **(J-N)**, suggesting that PLN-localized ferric iron-laden macrophages may derive in part from the afferent synovial tissue. Human synovial samples from subjects with osteoarthritis **(O)** compared to rheumatoid arthritis **(P)** demonstrated the clinical correlate of the Fe^3+^-laden macrophage phenotype. Statistics: Unpaired t-test **(C)**, linear regression **(D)**, and one-way ANOVA with Tukey’s multiple comparisons **(I**,**N)**; *\*p<0*.*05, **p<0*.*01, ***p<0*.*001*.

### Co-Stimulation of Alcam^+^ Macrophages and CD6^+^ T-Cells Potentially Promote IgG2b^+^ Plasma Cell Differentiation in Advanced TNF-Tg PLNs

Based on the findings of increased IgG2b^+^ plasma cells and Fe^3+^ macrophages localized to the Marco^+^ sinus region in Advanced PLNs, we evaluated changes in these cell populations by scRNAseq during Early and Advanced arthritis. Following integration of the Early (8192 cells) and Advanced (8136 cells) datasets, unsupervised clustering (resolution = 0.5) resolved 18 unique cell populations (Fig. 5A, complete annotations provided in Supplementary Table 3). Comparison of the cellular composition of Early versus Advanced PLNs revealed a striking increase in the number of T-cells (Fig. 5B, dashed black arrows; 2.0% Early vs 15.8% Advanced of clusters 5, 7, and 11 combined relative to total cells). Similarly, the monocyte / macrophage populations exhibited a notable increase in the Advanced condition (Fig. 5B, dashed grey arrows; 3.6% Early vs 7.4% Advanced of clusters 10, 12, 13, 16, and 18 combined relative to total cells). These proportional changes between the conditions are highlighted in Supplementary Figure 8 and Supplementary Table 3.

**Figure 5.**
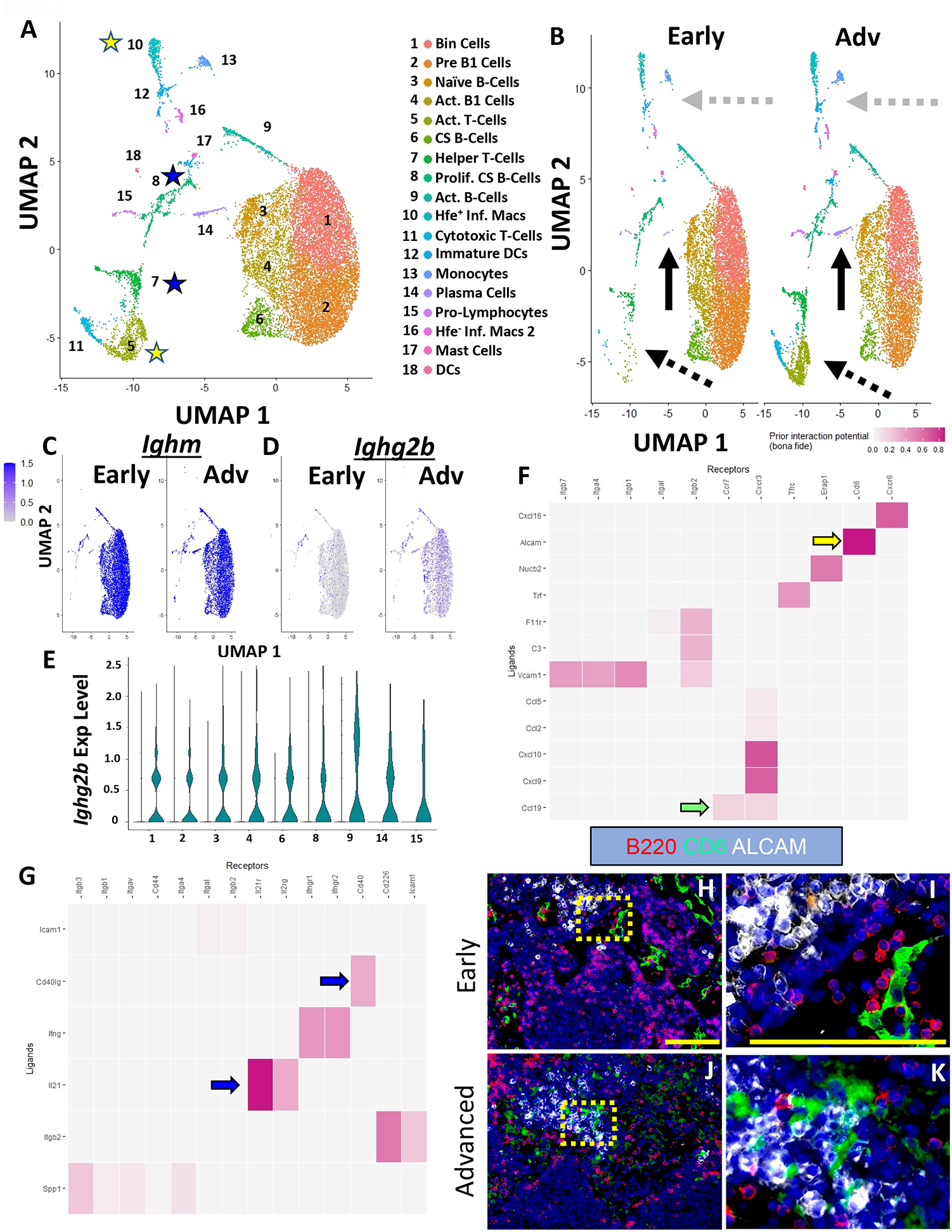
Co-Stimulation of Alcam^+^ Macrophages and CD6^+^ T-Cells Potentially Promote IgG2b^+^ Plasma Cell Differentiation in Advanced TNF-Tg PLNs. Single-cell sequencing resolved 18 distinct cell clusters **(A)**, representing subtypes of B-cells, T-cells, and monocytes / macrophages (annotations in Supplementary Table 3). Increased plasma cells (solid black arrows), T-cells (dashed black arrows), and macrophages (dashed grey arrows) were identified in Advanced PLNs **(B)**. Gene expression analysis revealed ubiquitous *Ighm* expression **(C)** in subset B-cell populations with a remarkable increase in *Ighg2b* expression **(D)** that was active across all B-cell subsets **(E)**. Cell-cell interaction analysis between *Hfe*^*+*^*/Cx3cr1*^*+*^*/Cd88*^*+*^*/Aif1*^*+*^ inflammatory macrophages (ligands) and *Ccr7*^*+*^*/Cd27*^*+*^*/Cd96*^*+*^*/Cd226*^*+*^ activated T-cells (receptors) (yellow stars in **A**) revealed high interaction potential between *Alcam*^*+*^ macrophages and *Cd6*^*+*^ T-cells (yellow arrow) with potential *Ccr7*/*Ccl19* (green arrow) recruitment mechanisms **(F)**. Similar analysis between *Cd4*^*+*^*/Cd8*^*-*^*/Cd40lg*^*+*^ activated helper T-cells (ligands) and *Nme1/2*^*+*^*/Mki67*^*+*^*/Top2a*^*+*^*/Jchain*^*+*^ proliferative class-switching B-cells (receptors) (blue stars as in **A**) showed high interaction potential with *Cd40lg* / *Il21* (T-cells) and *Cd40* /*Il21r* (B-cells) towards class-switching and plasma cell differentiation **(G**, blue arrows). Immunostaining further demonstrated the physical separation of Alcam^+^ and CD6^+^ cells in Early PLNs **(H**,**I)**, while these cells are in close proximity in Advanced arthritis **(J**,**K)**. Yellow scale bar = 100μm **(H-K)**.

The scRNAseq also further validated the spatial transcriptomics and immunofluorescent analysis with a remarkable increase in the proportion of *Cd93*^*+*^*/Irf4*^*+*^/*Cxcr4*^*+*^ plasma cells (Fig. 5B, solid black arrows; 0.08% Early vs 2.0% Advanced relative to total B-cells). Through investigation of immunoglobulin expression within the B-cell populations, we demonstrated the ubiquitous expression of *Ighm* within all B-cells (Fig. 5C). In addition, we confirmed the increase in *Ighg2b* levels in Advanced compared to Early TNF-Tg PLNs, but unexpectedly noted that without spatial differentiation, single-cell *Ighg2b* gene expression was elevated in all B-cell subsets and was not exclusive to differentiated plasma cells (Fig. 5D,E). However, the increase in *Ighg2b* expression in B-cells throughout the PLN was also specific relative to other immunoglobulins, where *Ighg1, Ighg2a/c, Ighg3*, and *Ighg4* showed limited or absent levels in both the Early and Advanced datasets (Supplementary Figure 9).

Given the robust validation of IgG2b^+^ class-switching with increased propensity towards plasma cell phenotype in the B-cells within Advanced PLNs (Figure 3), we next evaluated the potential for macrophage / T-cell stimulation leading to the differentiation of B-cells towards IgG2b class-switched plasma cells via T-cell help. We performed cell-cell interaction analysis of the clusters in the single-cell datasets using NicheNet R package (Browaeys et al., 2019).

Analysis of *Hfe*^*+*^/*Cx3cr1*^*+*^/*Cd88*^*+*^/*Aif1*^*+*^ inflammatory macrophages and *Ccr7*^*+*^/*Cd27*^*+*^/*Cd96*^*+*^/*Cd226*^*+*^ activated T -cells (yellow stars in Fig. 5A and yellow rows in Supplementary Table 3) showed a strong interaction potential between *Alcam* (macrophages) and *Cd6* (T-cells) (Fig. 5F, yellow arrow). Of note, Alcam / CD6 interaction is a co-stimulatory pathway involved in T-cell activation (Nair et al., 2010; Zimmerman et al., 2006), and anti-CD6 clinical trials for RA are currently underway with promising outcomes (Rodriguez et al., 2017; Rodriguez et al., 2012). Chemotactic signals, such as recruitment of *Ccr7*^+^ T-cells via *Ccl19*^+^ produced by macrophages, were also identified as a putative explanation for the dramatic increase in T-cells in Advanced arthritis (Figure 5F, green arrow). The activated *Cd4*^*+*^/*Cd8*^*-*^ /*Cd40lg*^*+*^ helper T-cells were also analyzed against the actively class-switching and proliferating *Nme1/2*^*+*^/*Mki67*^*+*^/*Top2a*^*+*^/*Jchain*^*+*^ B-cells (blue stars in Fig. 5A and blue rows in Supplementary Table 3). The interaction analysis identified well-described pathways including CD40LG (T-cell) and CD40 (B-cell) co-stimulation required for B-cell activation (Aversa et al., 1994; Durandy and Kracker, 2012), and IL21 (T-cell) and IL21R (B-cell) promoting plasma cell differentiation (Fig. 5G, blue arrow) (Moens and Tangye, 2014). Immunofluorescence of Early PLNs demonstrated a physical separation of CD6^+^ T-cells and Alcam^+^ macrophages (Fig. 5H,I), while these cells exhibited remarkably close proximity in the Advanced condition (Fig. J,K), suggesting increased cell-cell interaction potential as predicted by the bioinformatic analysis. Taken together, the findings of increased IgG2b^+^ plasma cells in TNF-Tg Advanced PLNs with Alcam/CD6 co-stimulatory signaling supports the theory that these events may be the nidus for autoimmunity and flare during chronic inflammatory arthritis.

## Discussion

B-cell translocation into sinuses is known to be integral to the pathogenesis of PLN “Collapse” and initiation of Advanced inflammatory-erosive arthritis in TNF-Tg mice (Bouta et al., 2018), but identification of distinct mechanisms involved in this process have been limited without adequate spatial resolution and analysis. Using multi-omic spatial and scRNA sequencing techniques, we identified dynamic changes in the cell populations and their transcriptomic profiles localized to Marco^+^ peri-follicular medullary sinuses of TNF-Tg PLNs during the progression of arthritis development. These alterations may be, in part, responsible for chronic inflammation induced autoimmunity and lymphatic dysfunction in RA. Previously, the transition from PLN expansion (Early arthritis) to PLN collapse (Advanced arthritis) was predominantly attributed to mechanical inhibition of fluid flow through B-cell accumulation in the sinuses (Bouta et al., 2018). However, the significant increase in IgG2b class-switching and accumulation of plasma cells within Advanced vs Early PLN sinuses suggests the possible onset of antigen-specific reactions and adaptive autoimmunity, which has never before been appreciated in the TNF-Tg mouse model of RA (Kontoyiannis et al., 1999; Li et al., 2010). This provides novel insights into previously reported B-cell effector functions in TNF-Tg mice, such as Bin cell translocation into PLN sinuses and identification of B-cells within the synovium and subchondral bone that promote erosion (Sun et al., 2018). Although the IgG2b class-switching was identified systemically, the lack of changes in serum IgG2b level was unexpected. Thus, future experiments are warranted to evaluate the antigen-specificity and B-cell receptor diversity associated with the IgG2b class-switching to formally evaluate the clonality of this humoral response. In addition, elucidating the mechanisms involved in the production of IgG2b^+^ cells are also required given the significant and positive correlation of PLN sinus *Ighg2b* expression with the severity of erosive arthritis in afferent ankles.

Moreover, the cellular dynamics in joint-draining lymph nodes in Advanced arthritis are far from limited to B-cells, but also exhibit substantial increases in macrophage and T-cell populations that likely regulate plasma cell differentiation. Through our spatial transcriptomics analysis, we identified down-regulation of Ferritin (*Fth1*) gene expression within the PLN sinuses of TNF-Tg mice with Advanced arthritis, which was associated with accumulation of Fe^3+^-laden macrophages. Our results also suggest strong cell-cell interaction between Alcam^+^ inflammatory macrophage expression and CD6^+^ recruited T-cells. Considering that the Alcam/CD6 co-stimulatory pathway has been implicated in collagen-induced arthritis and other models of autoimmunity (Fox, 2018; Li et al., 2020), and phase 1 clinical trials with anti-CD6 monoclonal antibody therapy in RA patients produced encouraging results (Rodriguez et al., 2017; Rodriguez et al., 2012), targeting this pathway to evaluate the effects on TNF-Tg arthritis and lymphatic pathology is warranted in future studies.

Most importantly, the current study also generates new hypotheses regarding the relationship between lymphatic dysfunction and arthritic flare, as illustrated in Figure 6, to guide future investigations. During Early arthritis, the B-cells within joint-draining PLNs begin exhibiting the first signs of activation with accumulation of CD21^+^/CD23^+^ Bin-cells as previously described (Kuzin et al., 2015; Kuzin et al., 2016; Li et al., 2010; Li et al., 2011; Moshkani et al., 2012). Then, the onset of Advanced arthritis can be caused by translocation of Bin cells with CD6^+^/CCR7^+^ T-cells into the Marco^+^ peri-follicular medullary sinuses where co-stimulation of Alcam^+^ macrophages/CD6^+^ T cells leads to IgG2b^+^ class-switching and plasma cell differentiation (Fig. 6B). Of note, in Early disease these macrophages, which could be egressing from inflamed joints, travel through the PLN with great speed in fully functioning lymphatics (Fig. 6C), and have no co-stimulatory effects on resident lymphocytes within the PLN. However, with the onset of Advanced arthritis and lymphatic dysfunction, these activated macrophages stagnate within the popliteal lymphatic vessels and PLN, and mediate tissue destruction and immune cell activation (Li et al., 2013). This lymphatic stagnation is associated with LEC (Bouta et al., 2017) and lymphatic muscle cell (LMC) (Jiao et al., 2022; Kenney et al., 2020; Kenney et al., 2022c; Liang et al., 2021) apoptosis and efferocytosis by joint-draining macrophages leading to intracellular iron accumulation (Fig. 6D). These iron-laden macrophages then likely exhibit reduced *Fth1* expression through negative feedback (Fig. 6E), passively flow into the efferent PLN, and exhibit an M1-polarized phenotype (DeRosa and Leftin, 2021) with secretion of chemokines (i.e., CCL19) to promote CCR7^+^ T-cell recruitment and B-cell translocation into the sinuses (Li et al., 2011). Within this microenvironment, we propose that Alcam co-stimulation of CD6^+^ T-cells promotes activation and differentiation of the recruited CCR7^+^ T-cells into CD4^+^ helper T-cells (Fig. 6F). These activated helper T-cells then interact with the Bin cells via CD40LG / IL21 (T-cells) and CD40 / IL21R (B-cells) (Fig. 6G) to promote IgG2b^+^ class-switch recombination and plasma cell differentiation (Fig. 6H). Without sufficient lymphatic flow, the IgG2b^+^ plasma cells then accumulate within the Marco^+^ peri-follicular medullary sinuses, but remain polyclonal in the context of sterile inflammation, and persist in the PLN without migration to the bone-marrow as long-lived plasma cells. Instead, these plasma cells physically inhibit passive lymphatic flow via clogging of the sinuses (Fig. 6I). In this double-hit hypothesis, with the loss of active lymphatic flow by LEC/LMC apoptosis and the inhibition of passive lymphatic flow by plasma cell clogging, the activated macrophages accumulate within the afferent synovium to promote onset of severe inflammatory-erosive arthritis (Fig. 6J). This revised model of lymphatic failure and arthritic flare provides new molecular and cellular targets for future experimentation, and potential insight into the pathogenesis of seronegative RA.

**Figure 6.**
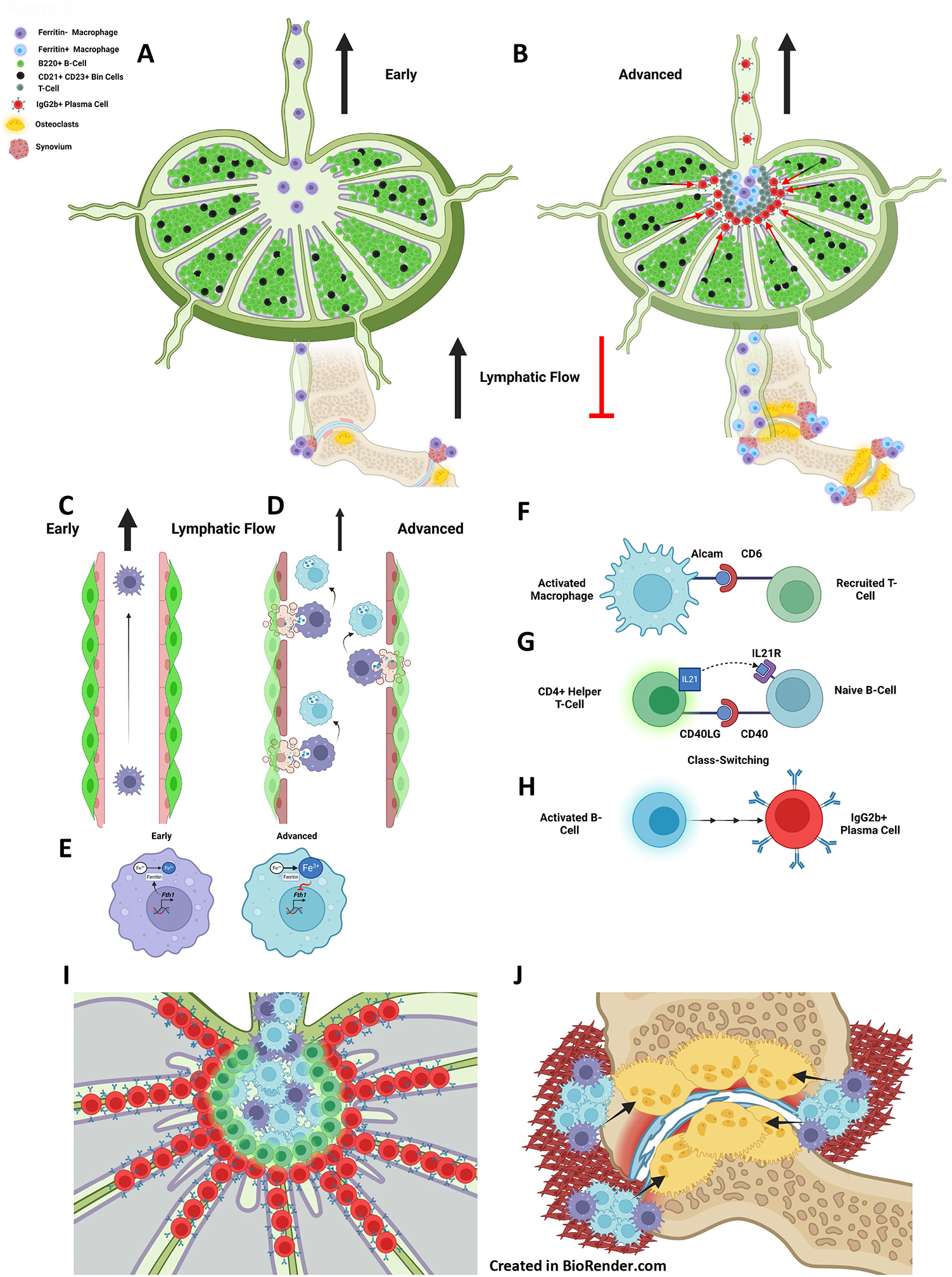
A new model of lymphatic failure and lymph node collapse with seronegative plasmocytic arthritic flare. We provided spatial and single-cell transcriptomic evidence that Bin cells (black), present in Early B-cell follicles (green) **(A)**, translocate into Marco^+^ perifollicular medullary sinuses in response to stimulation towards IgG2b class-switching and plasma cell differentiation during Advanced arthritis **(B**, red cells). Normal lymphatic drainage from Early RA joints **(C)** is lost in Advanced RA, associated with lymphatic muscle and endothelial cell death and macrophage efferocytosis, leading to Fe^3+^ accumulation as these cells attach to lymphatic vessels **(D)**. This inhibits further Ferritin production via negative feedback of *Fth1* gene expression **(E)**. The activated and Fe^3+^-laden macrophages stimulate the recruited T-cells into helper T-cells through interaction of the Alcam ligand (macrophage) and CD6 receptor (T-cell) **(F)**. These activated T-cells then stimulate the naïve Bin cells through CD40LG / IL21(T-cell) and CD40R / IL21R (B-cell) pathways **(G)**, which leads to IgG2b^+^ class-switching and plasma cell differentiation **(H)**. Passive lymphatic flow is then mechanically inhibited as the plasma cells accumulate and clog the sinuses **(I)**. The Fe^3+^-laden inflammatory macrophages then aggregate within the synovium of afferent joints to promote inflammation, osteoclastogenesis, bone resorption, and arthritic flare **(J)**.

To our knowledge, this effort marks the first attempt to elucidate lymphatic dysfunction during arthritic progression via spatial transcriptomics. However, our study has noteworthy limitations, including: 1) the inability of the approach to detect low level transcripts, protein synthesis, and post-translational modification, and 2) the absence of additional datasets from other RA animal models and human patients with Early and Advanced arthritis to corroborate our results. We also need to evaluate our molecular findings with lymphatic functional defects using the most current noninvasive near-infrared imaging methods (Proulx et al., 2017). Nevertheless, these novel findings demonstrate the power of spatial transcriptomics, in combination with scRNAseq, to address long-standing questions that could not be resolved by other approaches and encourages continued advances in the resolution of spatial transcriptomics technology from the tissue to the single-cell level.

## Supporting information

Supplementary Video Wild-Type

Supplementary Video Early TNF-Tg

Supplementary Video Advanced TNF-Tg

## Acknowledgements

We would like to thank the faculty and staff of the Histology, Biochemistry, and Molecular Imaging core, the Biomechanics and Multimodal Tissue Imaging core, and the Genomics Research Center at the University of Rochester for their contributions to this work. We would also like to thank the following funding sources: F30AG076326, T32GM007356, T32AR076950, R01AG059775, R01AR069000, R01AR056702, R00AR075899, OREF, Department of Medicine, R01AI111914, UH2AR067690, R21AR071670, and P30AR069655.

## Author Contributions

HMK, EMS, and CLW contributed to 1) the conception or design of the work, 2) acquisition, analysis, or interpretation of the data, and 3) drafted or revised the work. All other authors contributed to 1) the acquisition, analysis, or interpretation of the data and 2) drafted or revised the work. All authors have approved the submitted version of the manuscript. All authors also agree to be personally accountable for their contributions and ensure to appropriately respond to any questions related to the accuracy or integrity of the work.

