## Supplementary Figures for "Multi-omics Analysis Identifies IgG2b Class-Switching with ALCAM-CD6 Co-Stimulation in Lymph Nodes During Advanced Inflammatory-Erosive Arthritis"

[1]Center for Musculoskeletal Research; [2]Department of Pathology & Laboratory Medicine; [3]Department of Medicine, Division of Allergy, Immunology, Rheumatology; [4]Center for Vaccine Biology and Immunology; [5]Department of Microbiology and Immunology; [6]Genomics Research Center; [7]Department of Orthopaedics, [8]Department of Obstetrics and Gynecology, [9]Department of Neuroscience, [10]Department of Urology, University of Rochester Medical Center

**Supplementary Figure 1. Optimization of tissue permeabilization for PLNs.** PLNs from WT mice (n=2 mice, 4 PLNs) were harvested and processed to determine the optimal incubation for tissue permeabilization. A representative H&E-stained section is shown **(A)** along with representative images of TRITC-fluorescently tagged nucleotides after 3-minutes **(B)**, 6-minutes **(C)**, 12-minutes **(D)**, 18-minutes **(E)**, 24-minutes **(F)**, and 30-minutes **(G)** of tissue permeabilization. 12-minutes was the optimal time for permeabilization based on signal intensity and limited diffusion.

**Supplementary Figure 2. Integration of Advanced capture areas confirms consistency of the replicates.** We performed CCA integration of both Advanced capture areas and evaluated the consistency of the principal components of the two replicates embedded on a UMAP. Based on the direct overlay and lack of unique spot populations between Advanced 1 vs Advanced 2, we considered the Advanced capture areas to be replicates. Thus, the Advanced capture areas were merged together as a single group for comparison with the WT and Early capture areas.

**Supplementary Figure 3. PLN immunoglobulin expression and relationship with arthritic severity is selective for *Ighg2b*.** Corresponding with the immunoglobulin gene expression analysis and talus bone volumes in Figure 3, we also evaluated the levels of *Ighg1* and *Ighg2c* relative to *Ighm*. The expression of these genes is shown as a spatial feature plot overlaying H&E stained PLNs in the Early and Advanced condition **(A-F)**. In contrast to *Ighg2b* (Figure 3), neither *Ighg1* nor *Ighg2c* expression is increased in Advanced PLN sinuses, and the expression of these immunoglobulin genes does not correlate with talus bone volumes in the afferent ankles **(G-J)**.

**Supplementary Figure 4. IgM to IgG conversion with reduced B-cell proliferation in Advanced PLNs.** Relative to Early PLNs with abundant IgM$^+$ cells (red) **(A)**, Advanced PLNs exhibited a predominance of IgG$^+$ (green) cells **(B)**. Quantitatively, Early PLNs had significantly increased IgM$^+$ cells compared to Advanced **(C)**, while Advanced PLNs showed significantly increased IgG$^+$ cells **(D)**. The increased IgG$^+$ cells in Advanced PLNs was also associated with reduced cell proliferation measured by proliferating cell nuclear antigen (PCNA, red), which suggests that the IgG$^+$ cells represent mature non-proliferating plasma cells. Peanut agglutinin (PNA, green) marks B-cells that may associate with germinal centers **(E,F)**. Statistics: Unpaired t-test **(C,D)**, $**p<0.01$, $****p<0.0001$. Yellow scale bar = 100μm **(A,B,E,F)**.

**Supplementary Figure 5. Flow cytometry gating strategy for IgG2b$^+$ and CD138$^+$ cells.** Cells isolated from PLNs, spleen, blood, and bone marrow of TNF-Tg mice with Early and Advanced arthritis were processed for flow cytometry analysis to enumerate IgG2b$^+$ / CD138$^+$ cells. The Advanced PLN sample from Figure 3 is used as an example. First, cells were gated away from debris and remaining red blood cells by forward and side scatter **(A)**. Single cells were then selected on the linear diagonal of forward scatter height and area **(B)**. 7-AAD$^-$ (cell impermeable) live cells were then gated **(C)** and analyzed for surface expression of CD11b (macrophages) and CD3 (T-cells) **(D)**. All CD11b$^-$ / CD3$^-$ cells were considered to be predominantly B-cells and were used as the denominator to quantify the proportion of IgG2b$^+$ and CD138$^+$ cells **(E)**. The gating parameters for IgG2b and CD138 were validated by evaluating the gating on a CD11b$^-$ / CD3$^+$ T-cell population as a negative control for both markers **(F)**.

**Supplementary Figure 6. TNF-Tg mice with Advanced arthritis exhibit a systemic increase in IgG2b$^+$/CD138$^+$ plasma cells with no change in serum IgG isotypes.** Increased IgG2b$^+$/CD138$^+$ cells were also detected systemically in Advanced bone marrow **(A,B)**, blood **(C,D)**, and spleen **(E,F)**. As shown by the quantifications provided in Supplementary Table 2, the increase in IgG2b$^+$/CD138$^+$ was primarily driven by enhanced IgG2b expression in the spleen, blood, and bone marrow (no change in CD138 alone), while the PLNs were unique with a significant increase in both single-positive IgG2b and CD138 cell types ($p<0.05$). Interestingly, despite the increase in IgG2b$^+$/CD138$^+$ plasmablasts / plasma cells systemically, there was no difference in serum IgG2b antibody levels between Early and Advanced TNF-Tg mice. Both TNF-Tg cohorts exhibited expected proportions of immunoglobulin isotypes with IgG2b as the predominant isotype in C57BL/6 mice (Klein-Schneegans et al., 1989) **(G)**. Together, these data suggest that increases in CD138$^+$ plasmablasts / plasma cells are specific to the lymph nodes despite systemic increases in IgG2b-expression. Statistics: 2-way ANOVA with Tukey's multiple comparisons **(G)**, $^{**}p<0.01$.

**Supplementary Figure 7. Alcam$^+$ and F4/80$^+$ macrophages localize to Marco$^+$/Lyve1$^+$ lymphatic endothelial cell lined sinuses.** In a representative Advanced PLN, immunofluorescence revealed Alcam$^+$ (white) / F4/80$^+$ (red) macrophages closely localized to the Marco$^+$ (green) PLN sinuses **(A)**. Lyve1 (green) immunostaining was demonstrated in the same cellular pattern and location within the PLN sinuses as the Marco$^+$ cells in **A**, further confirming the Marco$^+$ lymphatic endothelial cell identity. On the other hand, peripheral node addressin (PNAd, red) marks high endothelial venules **(B)**. Note, as the F4/80$^+$ cells localized to similar regions of the PLN as the iron-laden cells, and because macrophages are known to be the primary immune cell regulator of iron homeostasis (DeRosa and Leftin, 2021), the iron-laden cells are presumed to be macrophages. Yellow scale bar = 100μm **(A,B)**.

**Supplementary Figure 8. Increased proportion of T-cells and monocyte / macrophage populations in TNF-Tg PLNs with Advanced arthritis.** The 18 cell clusters identified by single-cell RNA-sequencing in Figure 5 were merged to highlight the general B-cell, T-cell, and monocyte / macrophage cell types. Comparison of Early versus Advanced **(A)** PLNs demonstrated a dramatic increase in the number of T-cells (green; 2.0% Early **(B)** vs 15.8% Advanced **(C)**). A similar increase was notable in the proportion of monocytes / macrophages in Advanced PLNs (3.6% Early **(B)** vs 7.4% Advanced **(C)**). In combination with Figure 5, these findings suggest that increased T-cells and monocytes / macrophages are accumulating in Marco$^+$ sinuses to ultimately stimulate IgG2b$^+$ class-switching and plasma cell differentiation of intra-sinus B-cells of TNF-Tg PLNs.

**Supplementary Figure 9. B-cell immunoglobulin expression is selective for *Ighg2b* in Advanced PLNs.** To validate the increased *Ighg2b* expression noted within the PLNs of TNF-Tg mice with Advanced arthritis by spatial transcriptomics, we evaluated the immunoglobulin isotypes between the Early and Advanced conditions after subsetting the B-cell populations by scRNAseq. As shown in Figure 5, there was high gene expression of *Ighm* within all B-cell subtypes, while Advanced B-cells showed increased *Ighg2b* levels compared to Early. The increased immunoglobulin expression in Advanced B-cells was selective to the *Ighg2b* isotype with limited expression in both Early and Advanced conditions for *Ighg1* **(A)**, *Ighg2c* **(B)**, and *Ighg3* **(C)**. *Ighg2a* and *Ighg4* were undetectable throughout the datasets, and are thus omitted.

**Supplementary Table 1. RIN values for estimating RNA integrity.** Prior to utilization of the PLN tissue blocks for analysis, ten-consecutive 10µm sections were collected to evaluate RNA quality through RIN quantification. We considered RIN values >8.0 to be optimal for gene expression analysis, and all tissue showed RIN values in this range with an overall minimum RIN value of 9.4 (Optimization). There was a minimum RIN value of 9.6 for the gene expression blocks (all TNF-Tg), and a maximum RIN value of 10.0 (WT), which is the highest possible RIN value.

**Supplementary Table 2. Quantification of IgG2b$^+$ and CD138$^+$ cells in the bone marrow, blood, spleen, and PLN of TNF-Tg cohorts by flow cytometry.** To assess the numbers of IgG2b$^+$ and/or CD138$^+$ cells in TNF-Tg mice with Early or Advanced arthritis, we harvested cells from the bone marrow, blood, spleen, and PLNs for flow cytometry (n = 3 mice per group). Based on the results described in Figure 3, Supplementary Figure 6, and the gating strategy in Supplementary Figure 5, the mean ± standard deviation for the proportion of IgG2b$^+$ and/or CD138$^+$ cells relative to total CD11b$^-$/CD3$^-$ cells is provided. While all organs showed increased IgG2b$^+$ and IgG2b$^+$/CD138$^+$ cells in Advanced vs Early TNF-Tg mice, the PLNs were unique to all other compartments with a significant increase in CD138$^+$ cells alone ($p<0.05$). Thus, the increase in IgG2b$^+$/ CD138$^+$ cells in the bone marrow, blood (not significant), and spleen were primarily driven by increases in IgG2b$^+$ cells, while both IgG2b and CD138 were enhanced in the PLNs of TNF-Tg mice with Advanced arthritis.

**Supplementary Table 3. Single-cell cluster identity.** Corresponding with the 18 clusters shown by UMAP (Figure 5), the complete cluster identities and proportion of total cells for the Early and Advanced conditions are provided in this table. The complete cluster identities provide the associated genes and annotations of each cluster number.

**Supplementary Videos.** The videos demonstrate a fly-through of an image-stack from the original Prussian Blue and Nuclear Fast Red stained images. Note the rotation artifact of the images, which is a product of the alignment procedures performed in Amira software. Color segmentation was performed in ImageJ using the Color Deconvolution 2 plug-in with a threshold set on the Feulgen LightGreen vector. The resultant segmentations were imported into Amira, aligned with the histology, and rendered in 3D to depict the lymph node stromal cells (red/orange) and the iron-laden cells within the sinuses (blue). The video animations were created in the Amira software.

Supplementary Figure 1

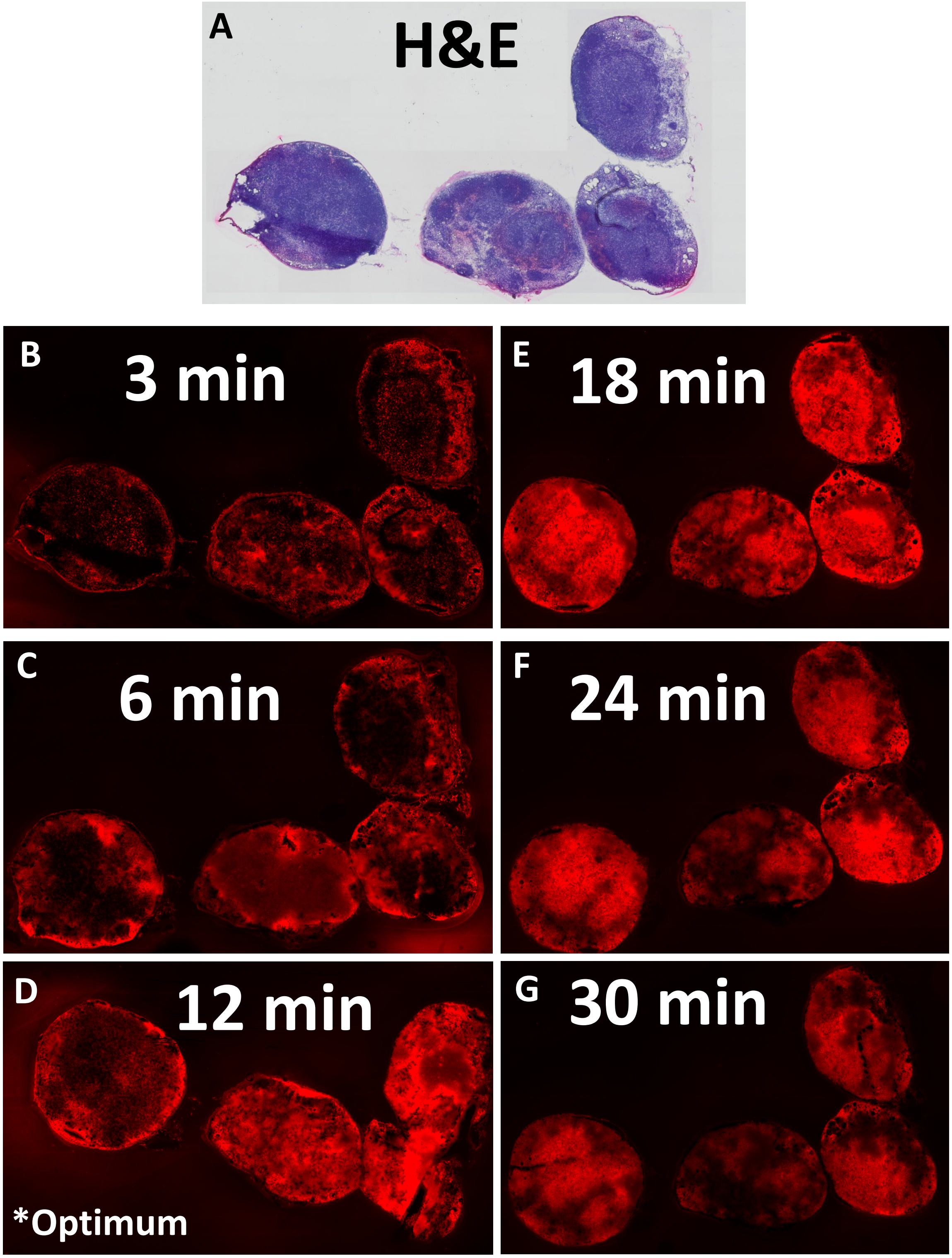

Supplementary Figure 2

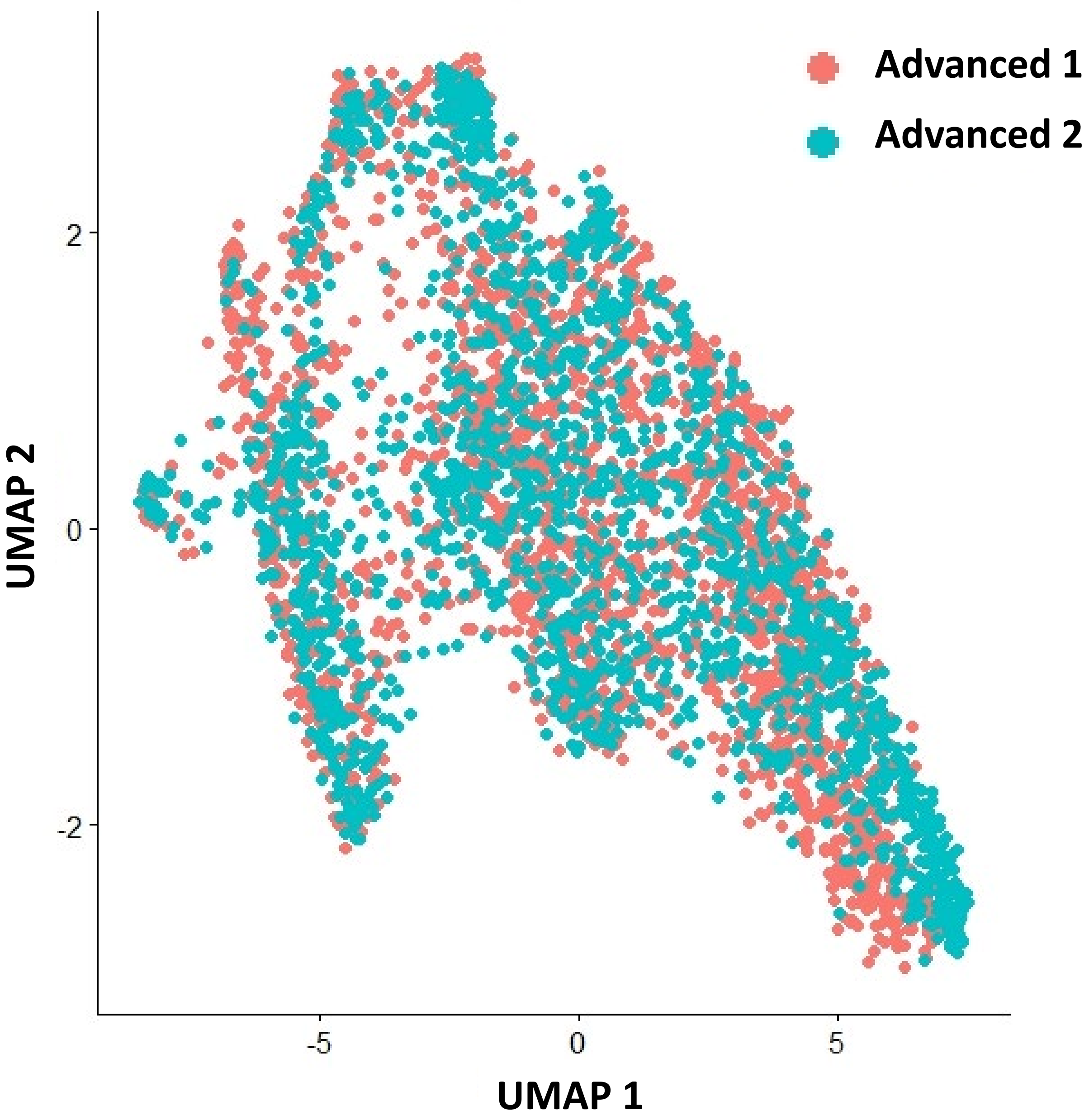

Supplementary Figure 3

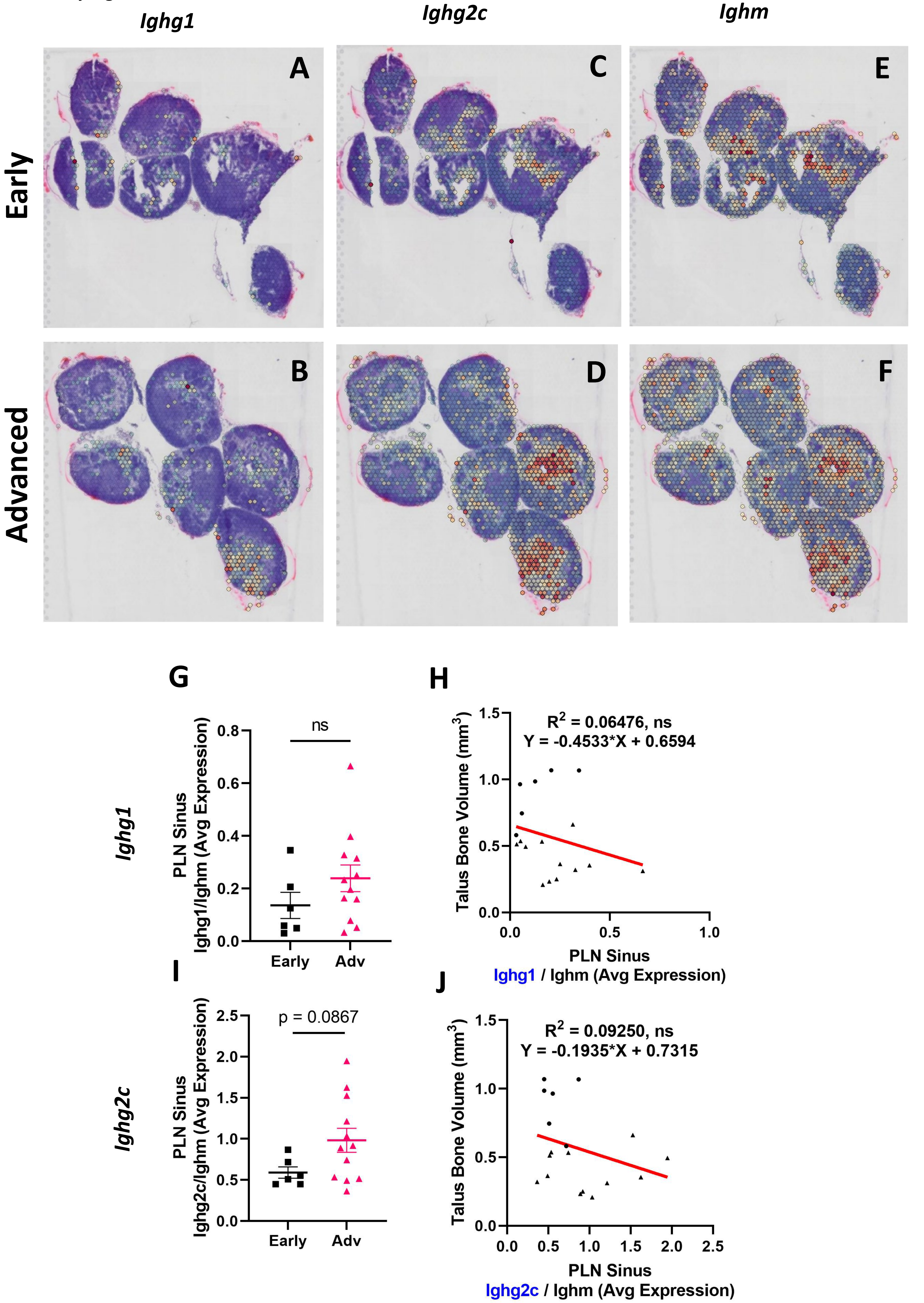

Supplementary Figure 4

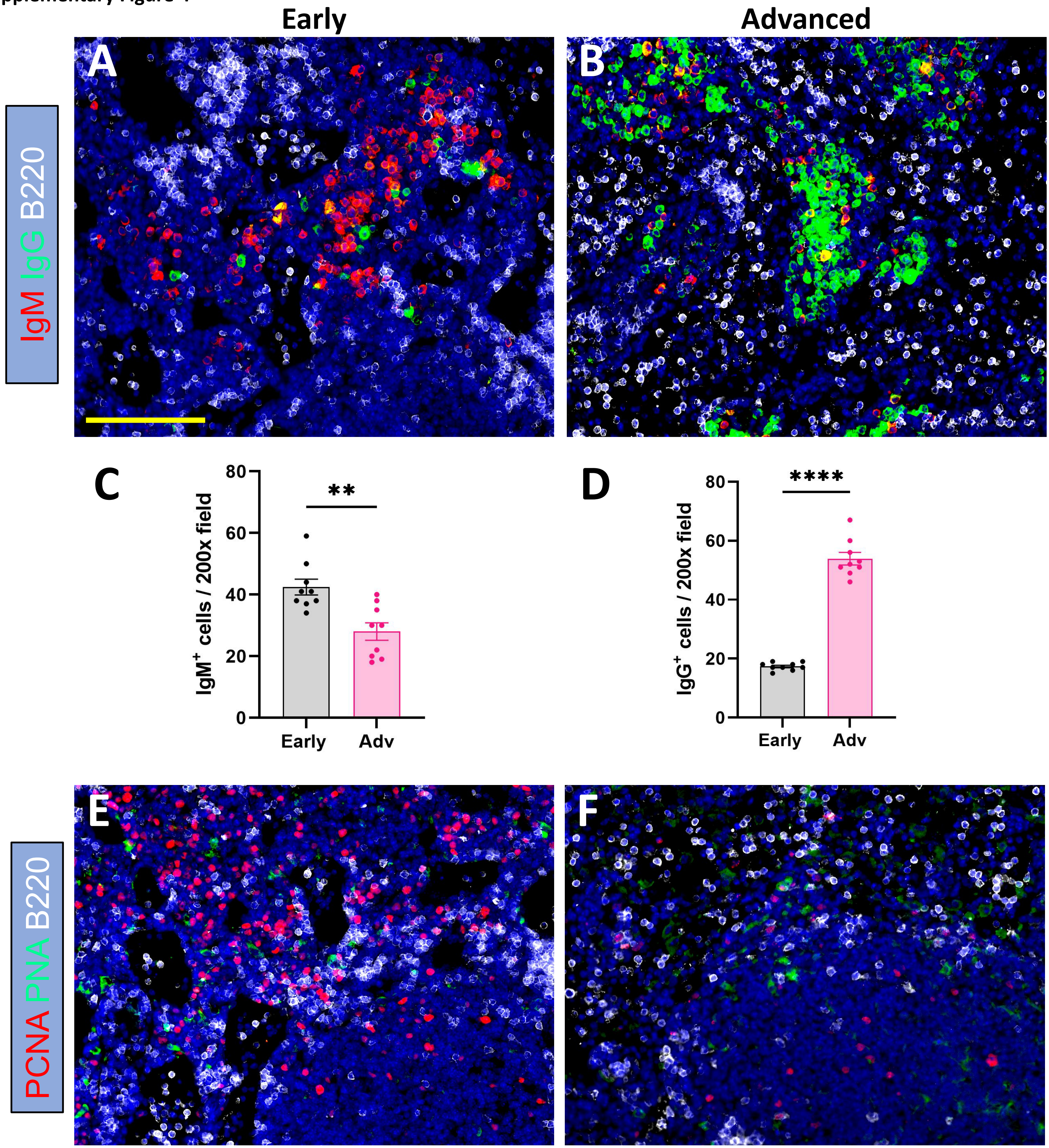

Supplementary Figure 5

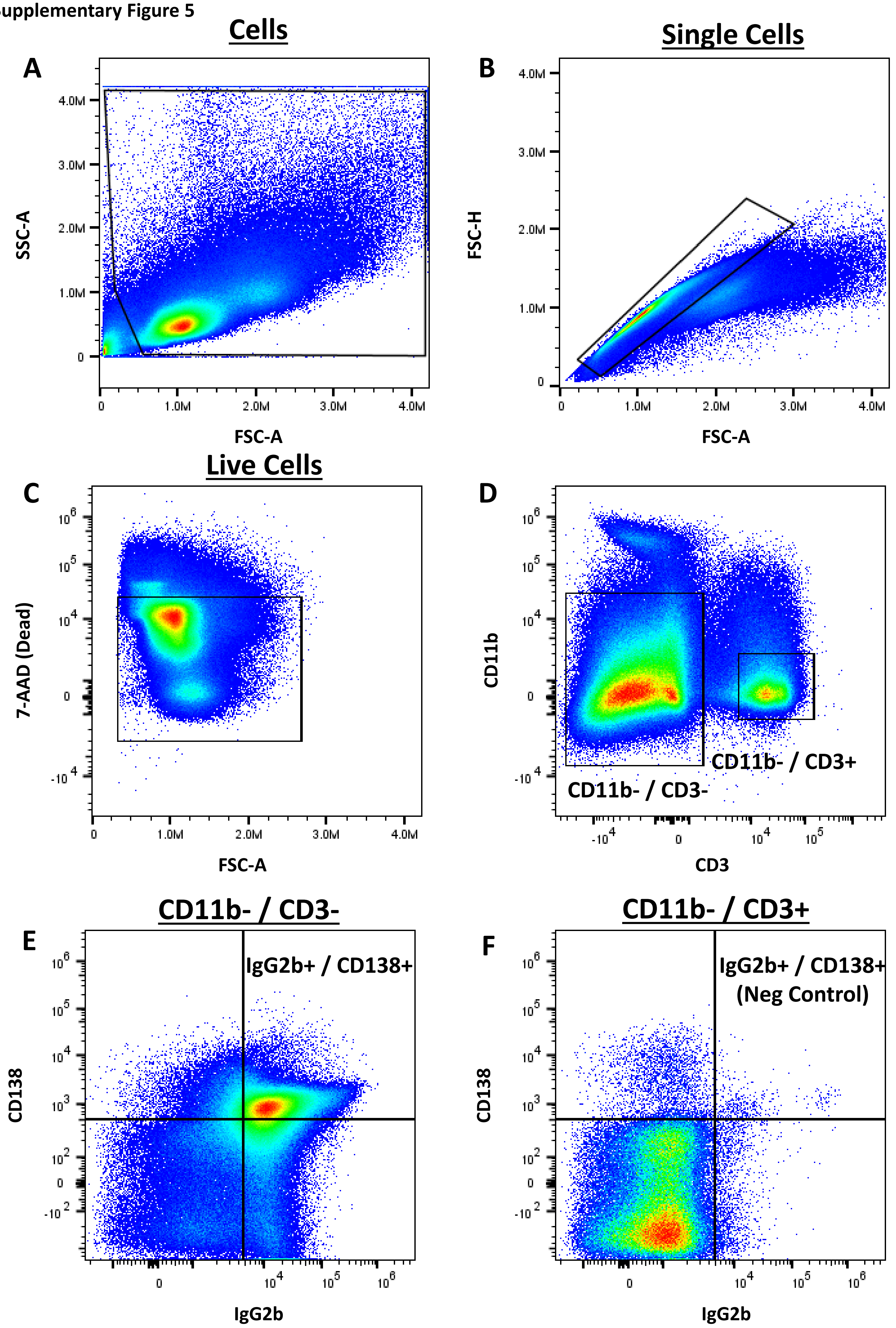

Supplementary Figure 6

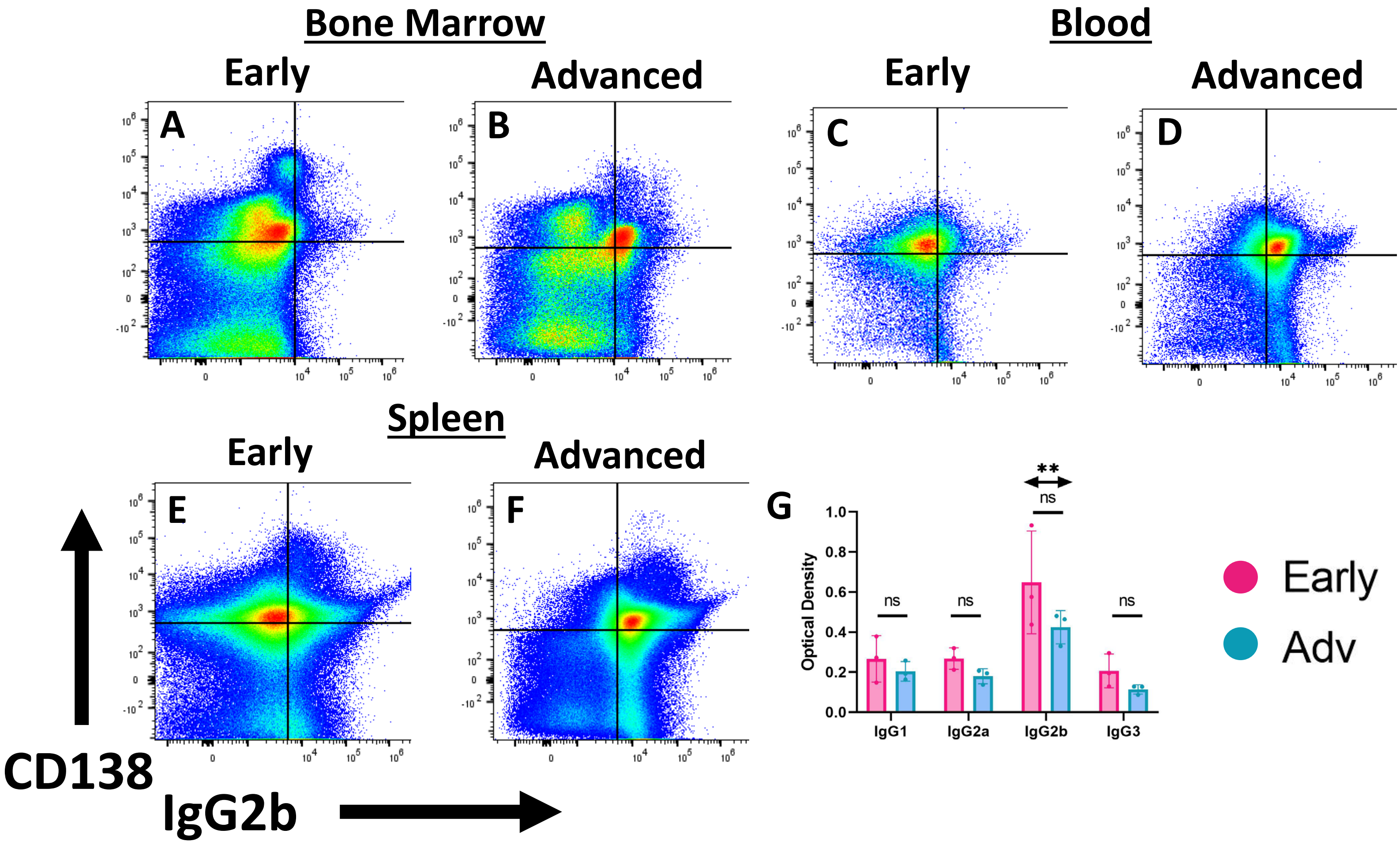

**Supplementary Figure 7**

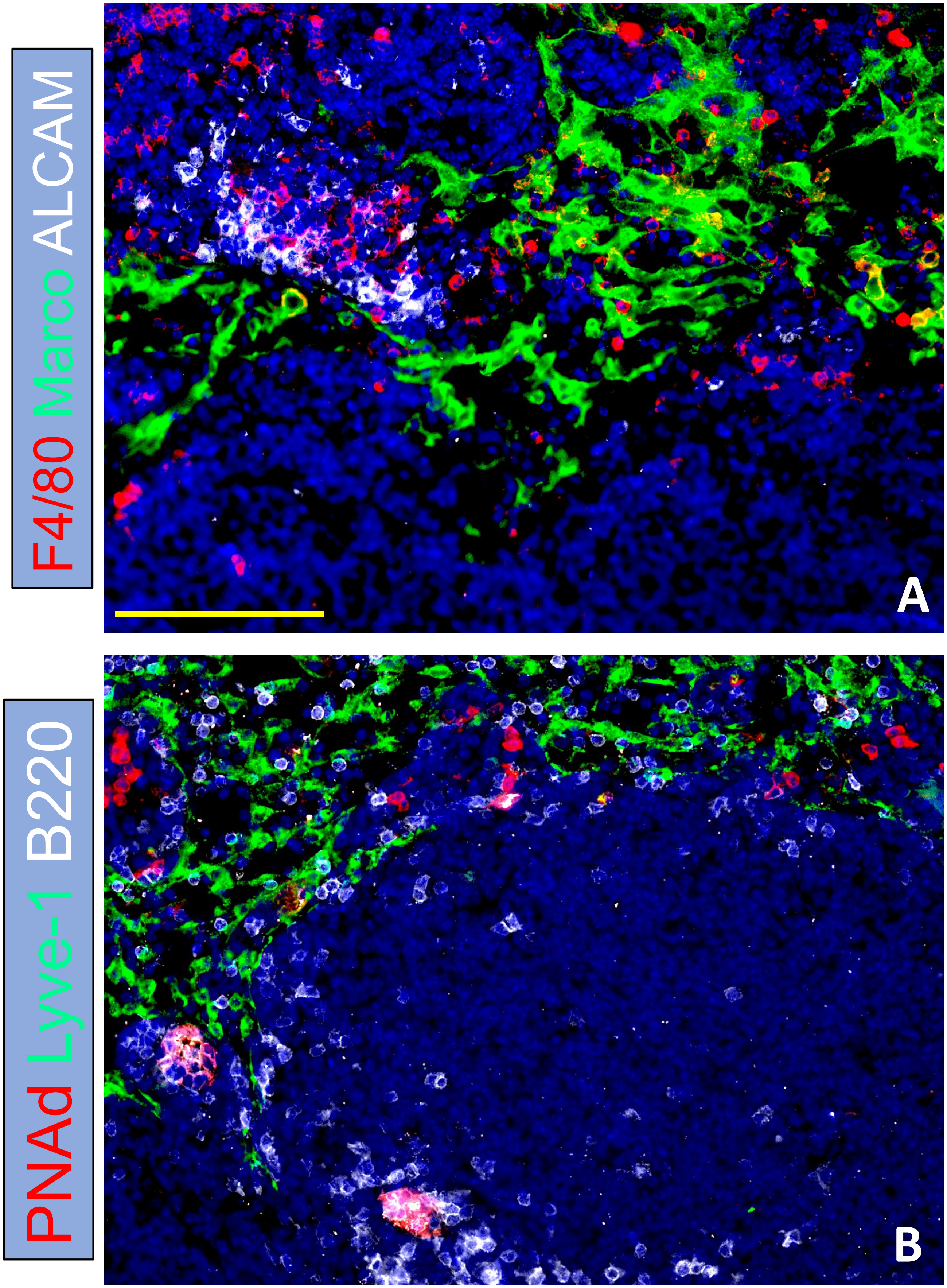

Supplementary Figure 8

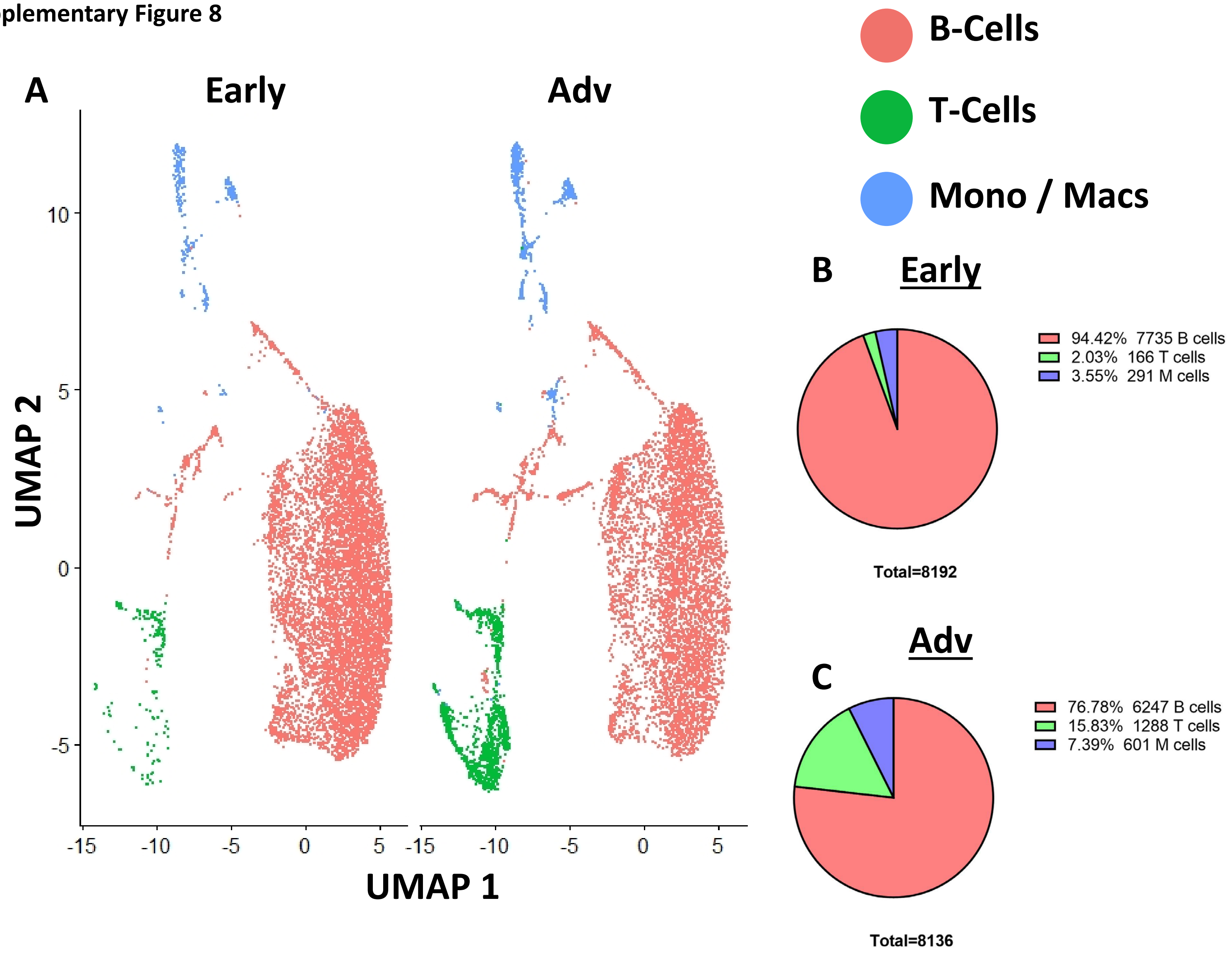

Supplementary Figure 9

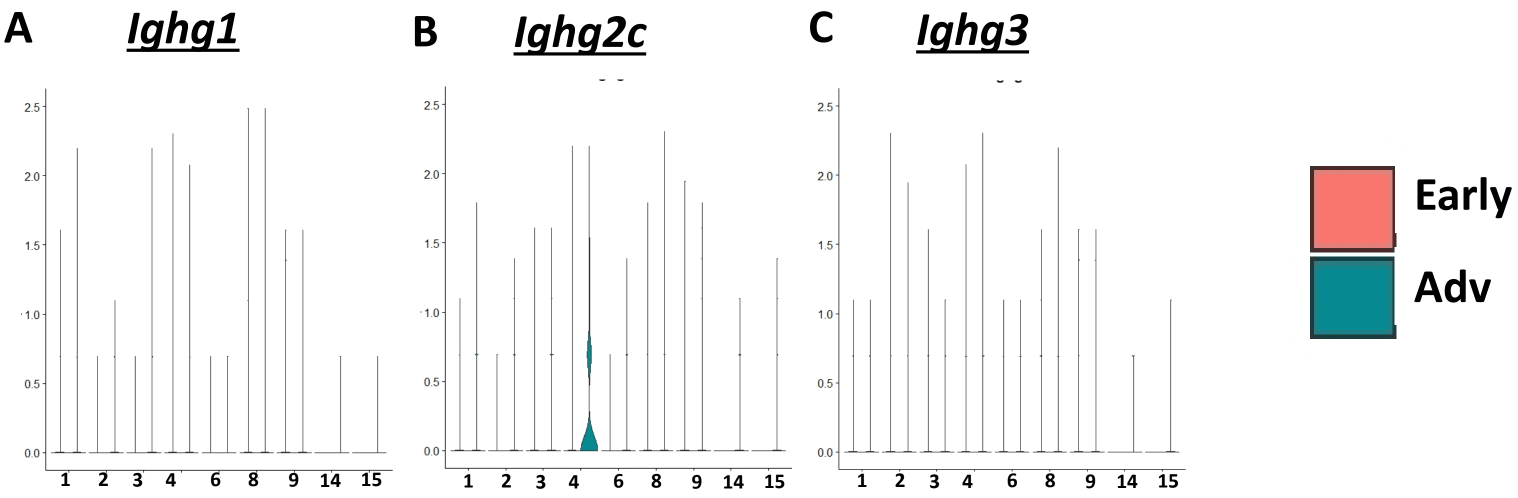

**Supplementary Table 1**

| **Sample** | **RNA Integrity Numbers (RIN)** |
|---|---|
| Optimization | 9.4 |
| Wild-Type | 10.0 |
| TNF-Tg Early | 9.6 |
| TNF-Tg Advanced 1 | 9.6 |
| TNF-Tg Advanced 2 | 9.6 |

**Supplementary Table 2**

| | IgG2b+ | | | CD138+ | | | IgG2b+ / CD138+ | | |
|---|---|---|---|---|---|---|---|---|---|
| **Organ** | **Early** | **Adv** | ***p*-value (t-test)** | **Early** | **Adv** | ***p*-value (t-test)** | **Early** | **Adv** | ***p*-value (t-test)** |
| **BM** | 4.2 ± 0.4% | 26.4 ± 9.6% | * 0.02 | 40.9 ± 3.7% | 30.1 ± 6.4% | ns, 0.07 | 2.7 ± 0.5% | 15.6 ± 6.6% | * 0.03 |
| **Blood** | 20.61 ± 8.5% | 54.6 ± 20.3% | ns, 0.06 | 61.1 ± 5.7% | 52.3 ± 12.4% | ns, 0.33 | 13.1 ± 5.8% | 35.1 ± 16.7% | ns, 0.10 |
| **Spleen** | 29.2 ± 6.3% | 66.3 ± 17.2% | * 0.02 | 50.0 ± 10.2% | 50.6 ± 16.2% | ns, 1.0 | 15.5 ± 5.9% | 42.13 ± 13.3% | * 0.03 |
| **PLN** | 28.89 ± 0.6% | 74.6 ± 7.6% | *** 0.0005 | 41.4 ± 12.1% | 66.9 ± 5.0% | * 0.03 | 11.4 ± 3.1% | 55.6 ± 4.6% | *** 0.0002 |

Supplementary Table 3

| Cluster # | Cluster Identity | Early (%) | Advanced (%) |
|---|---|---|---|
| 1 | CD55+ / Ighd+ / Cd21+ / Cd23 + Bin Cells | 32.45% | 26.62% |
| 2 | Ly6d+ / Mzb1+ Pre B1 Cells | 34.90% | 22.85% |
| 3 | Ighd+ / Vpreb3+ Naïve B-Cells | 9.13% | 9.08% |
| 4 | Ahr+ / Zbtb20+ / Mzb1+ / CD24a+ Activated B1 Cells | 9.11% | 6.00% |
| 5 | Ccr7+ / Cd27+ / Cd96+ / Cd226+ Activated T-Cells | 0.50% | 8.60% |
| 6 | Nme1/2+ Class-Switching B-Cells | 4.90% | 3.44% |
| 7 | Cd4+ / Cd8- / Cd40lg+ Activated Helper T-Cells | 1.32% | 4.14% |
| 8 | Nme1/2+ / Mki67+ / Top2a+ / Jchain+ Proliferative Class-Switching B-Cells | 2.08% | 2.97% |
| 9 | Sec61a1+ Activated B-Cells | 1.65% | 3.18% |
| 10 | Hfe+ / Cx3cr1+ / Cd88+ / Aif1+ Inflammatory Macrophages | 1.54% | 2.56% |
| 11 | Cd4- / Cd8+ / Ly6c2+ / Cd160+ / Cxcr3+ Activated Cytotoxic T-Cells | 0.21% | 3.09% |
| 12 | Tmem176a+ / Cx3cr1+ Immature Dendritic Cells | 0.77% | 2.36% |
| 13 | Ly6c2+ / Ccr2+ / Itgax+ Monocytes | 0.74% | 1.51% |
| **14** | **Cd93+ / Irf4+ / Cxcr4+ Plasma Cells** | **0.07%** | **1.52%** |
| 15 | Dntt+ / Sox4+ Pro-Lymphocytes | 0.15% | 1.11% |
| 16 | Hfe- / Cd86+ / Vcam1+ / Aif1+ Inflammatory Macrophages | 0.34% | 0.71% |
| 17 | Kit+ / Mcpt4+ / Cma1+ Mast Cells | 0.35% | 0.58% |
| 18 | Cd207+ / Cd8+ Dendritic Cells | 0.16% | 0.25% |
